# Wnt5a gain- and loss-of-function present distinctly in craniofacial bone

**DOI:** 10.1101/2025.07.21.665966

**Authors:** Claire J. Houchen, Portia Hahn Leat, Cassandra Delich, Jocelyn Vang, Sara Haggard, Joseph L. Roberts, Hicham Drissi, Erin E. Bumann

## Abstract

**Introduction:** Robinow syndrome has characteristic craniofacial and dental features and can be caused by gain- or loss-of-function variants in Wnt family member 5A (*WNT5A*) non-canonical signaling. The craniofacial and dental manifestation of Robinow syndrome is heterogenous, as is the effect of altered *Wnt5a* in animal models. The relationship between *Wnt5a* and craniofacial and dental phenotypes is not fully understood.

**Methods:** To investigate the role of *Wnt5a* in craniofacial and dental development, we utilized a *Wnt5a* conditional loss-of-function (LOF: *Wnt5a*^*fl/fl*^*;Ctsk*^*cre*^) and a *Wnt5a* conditional gain-of-function (GOF: *Rosa26-LSL-Wnt5a;Ctsk*^*cre*^) model to determine the effect of both LOF and GOF of *Wnt5a* in bone cells during craniofacial and dental development. Postnatal day 10 conditional LOF *Wnt5a*, GOF *Wnt5a*, and control skulls were scanned by micro-computed tomography and assessed using traditional and geometric morphometrics. Mandibular bone apoptosis was further assessed by TUNEL staining.

**Results:** Conditional *Wnt5a* LOF resulted in midface hypoplasia, increased maxillary intermolar width, increased rostral basisphenoid width, and delayed molar eruption. *Wnt5a* LOF mandibles did not have altered bone mineral density or bone microarchitecture unlike our previous study examining *Wnt5a* LOF femurs. In contrast, conditional *Wnt5a* GOF results in macrocephaly, shortened hard palate, increased zygomatic length, micrognathia, and mandibular process morphology changes. The micrognathia and mandibular process morphology changes in the *Wnt5a* GOF mice were not due to increased apoptosis. A partially penetrant snout deviation was present in both the *Wnt5a* LOF and GOF mice.

**Conclusions:** Craniofacial and dental phenotype differed between mice with conditional GOF and LOF of *Wnt5a*, consistent with the craniofacial phenotype heterogeneity in *Wnt5a*-associated Robinow syndrome. We detected tooth eruption delay, mandibular condyle dysmorphology, and facial asymmetry in mice with altered *Wnt5a* that have not been previously reported in patients. Our data suggest precise regulation of *Wnt5a* is essential for proper craniofacial and dental development.

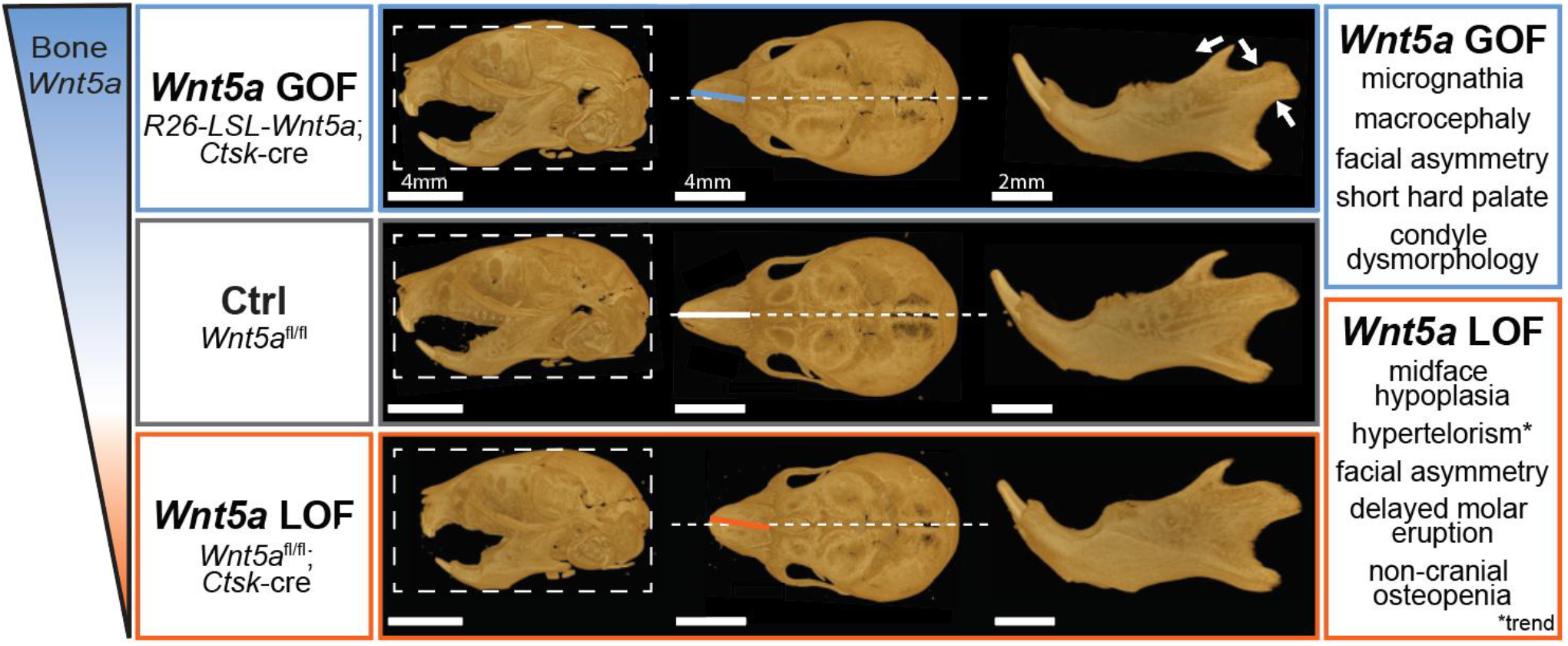

## INTRODUCTION

Craniofacial anomalies affecting the head, mouth, or jaw are common and affect 2-3% of all babies (Mossey and Catilla 2003). Craniofacial anomalies can be isolated and of non-syndromic origin or can occur as part of a syndrome. One such syndrome, Robinow syndrome, has characteristic craniofacial, oral, and dental differences (Bain et al. 1986; Wadlington et al. 1973). Robinow syndrome occurs due to pathogenic variants in Wnt family member 5A (*WNT5A)* or other members of the *WNT5A* non-canonical signaling pathway (Person et al. 2010; Roifman et al. 2015). There is heterogeneity in the specific craniofacial manifestations of Robinow syndrome, even among individuals with mutations in the same gene. For example, the craniofacial phenotype of individuals with *WNT5A*-associated Robinow syndrome include midface hypoplasia, hypertelorism, macrocephaly, short hard palate, micrognathia, cleft lip/palate, and others, but not all pathogenic variants in *WNT5A* result in the same set of craniofacial anomalies (Conlon et al. 2021; White et al. 2018). Unfortunately, the rarity and heterogeneity of Robinow syndrome makes affected individuals challenging to identify for referral for proper diagnostic testing (Conlon et al. 2021). A greater understanding of how Robinow syndrome dentally and craniofacially manifests is critical for decreasing time to diagnosis and improving treatment for affected individuals.

Recent evidence suggests that perturbations in *WNT5A* non-canonical signaling from loss-of-function, gain-of-function, or hypomorphic variants in *WNT5A* can all result in Robinow syndrome (White et al. 2018; Zhang et al. 2022; Zhang et al. 2021). The relationship between variant effect and craniofacial phenotype is not fully understood, but animal models have begun to shed light on the role of *Wnt5a* in craniofacial and dental development (Hosseini-Farahabadi et al. 2013; Hosseini-Farahabadi et al. 2017; Lin et al. 2011; Tokavanich et al. 2021; Yamaguchi et al. 1999). Severe skeletal defects including an extreme craniofacial phenotype can be observed in E18.5 *Wnt5a*^-/-^ embryos, but *Wnt5a* global knockout is perinatal lethal in mice and is consequently neither usable for assessing postnatal craniofacial morphology nor likely representative of the Robinow syndrome patient population (Yamaguchi et al. 1999). Dentally, *Wnt5a* is known to influence molar cusp morphology and root development, and is robustly expressed in alveolar bone during tooth eruption, however, whether alterations in non-canonical Wnt signaling results in eruption changes is not fully understood (Lin et al. 2011; Sarkar and Sharpe 1999; Tokavanich et al. 2021; Xiang et al. 2014). This study compares the craniofacial and dental phenotype of mice with altered *Wnt5a* using a cathepsin K cre *(Ctsk*^*cre*^) model to determine the extent to which craniofacial phenotypes differ by *Wnt5a* function in bone and fibrocartilage (Fig 1A-C).

**Figure 1.**
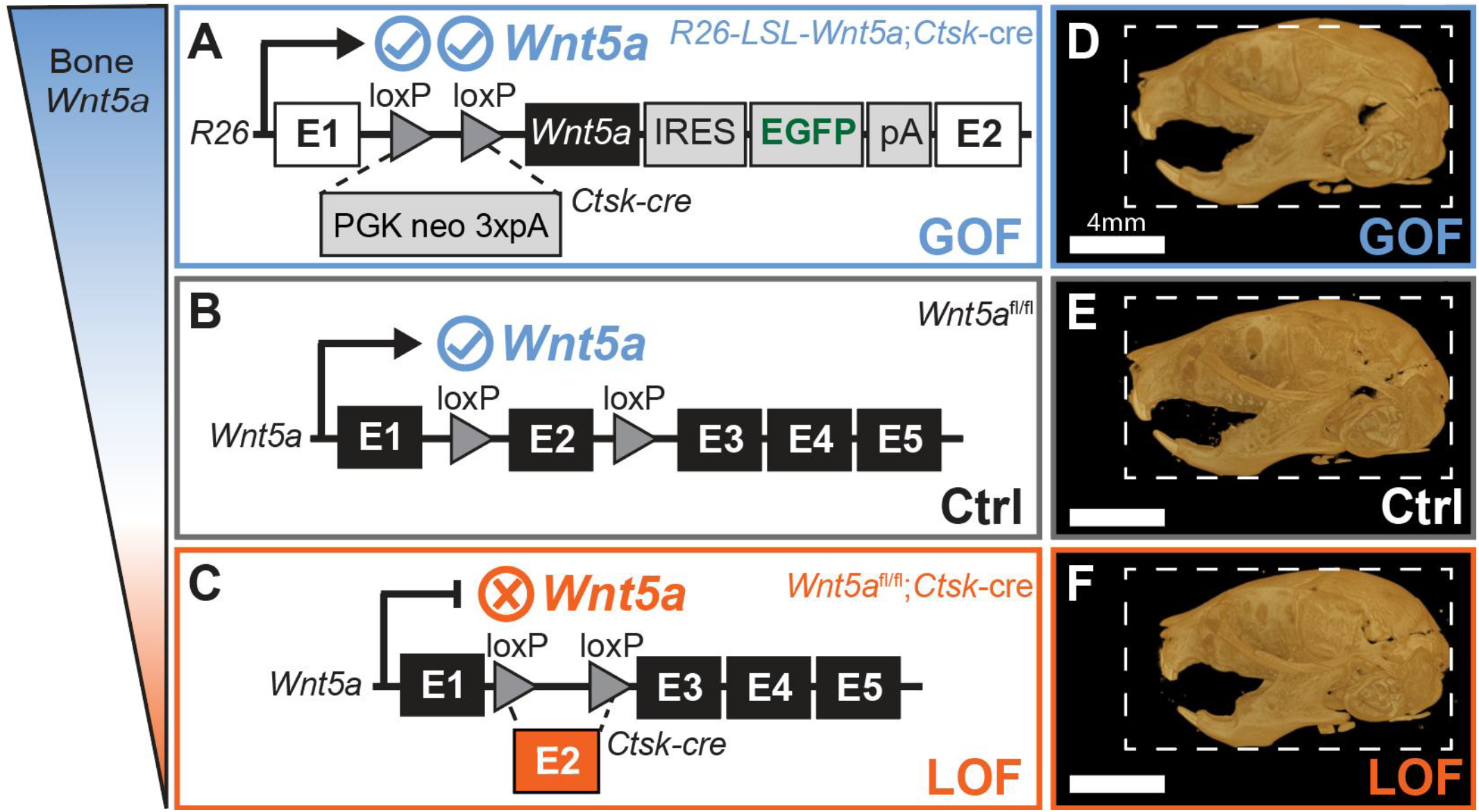
*Wnt5a* gain-of-function (GOF), control (Crtl), and loss-of-function (LOF) mice. **A)** Conditional *Wnt5a* GOF (*Rosa26-LSL-Wnt5a;Ctsk*^*cre*^) mice (Chen et al. 2014), **B)** control *Wnt5a* floxed mice (*Wnt5a*^*fl/fl*^), and **C)** conditional *Wnt5a* LOF(*Wnt5a*^*fl/fl*^*;Ctsk*^*cre*^) genotype (Roberts et al. 2020) descriptions. Lateral micro-CTs of postnatal day 10 **D)** *Wnt5a* GOF, **E)** control, and **F)** *Wnt5a* LOF mice demonstrate the craniofacial and dental phenotype of each group. Dashed line indicates control skull size.

Originally the *Ctsk*^*cre*^ model was designed to investigate bone-resorbing osteoclasts, but the cre recombinase is also expressed bone-depositing osteoblasts, osteocytes, bone lining cells in bone, and fibrocartilage, in addition to other organs, tissues, and cell types (Chai et al. 2023). In the mandible of *Ctsk*^*cre*^ mice, *cre* expression is in osteoclasts, periosteal cells, pericytes, bone marrow stromal cells and the fibrocartilage in the temporomandibular joint (TMJ) (Ding et al. 2022). Cathepsin K-positive cells express cre recombinase in the *Ctsk*^*cre*^ model after embryonic day 14.5 when cathepsin K expression can first be detected (Debnath et al. 2018). Cathepsin K-positive cells are essential for proper craniofacial and dental development, demonstrated by the craniofacial and dental changes in *Ctsk*^cre^;*DTA*^*fl/+*^ mice (Hassan et al. 2022).

In summary of the existing literature, both *Ctsk*^*cre*^ -expressing cells and *Wnt5a* are known to impact craniofacial and dental development. While the bone phenotype of *Wnt5a*^*fl/fl*^*;Ctsk*^*cre*^ mice has been examined in the appendicular (femur) and axial (vertebral body) skeleton (Roberts et al. 2020), the role of *Wnt5a* in *Ctsk*^*cre*^ -expressing cells is not described in craniofacial bone. To investigate the role of *Wnt5a* in craniofacial and dental development in postnatal embryos, we utilized a *Wnt5a* conditional knockout model (LOF: *Wnt5a*^*fl/fl*^*;Ctsk*^*cre*^) and a *Wnt5a* conditional overexpression model (GOF: *Rosa26-LSL-Wnt5a;Ctsk*^*cre*^) that allows us to investigate both LOF and GOF *Wnt5a* effects in bone cells on craniofacial development (Fig 1A-C). Outside of the Robinow syndrome patient population, this study demonstrates the important craniofacial and dental role of *Wnt5a*.

## MATERIALS AND METHODS

### Animals

Conditional *Wnt5a* knockout (LOF) mice were generated as previously reported (Roberts et al. 2020). Homozygous *Rosa26-LSL-Wnt5a* mice were crossed with homozygous *Ctsk*^*cre*^ to generate condition *Wnt5a* knock-in (GOF) mice. Mice were maintained in accordance with applicable state and federal guidelines and all experimental procedures were approved by the Atlanta VA Medical Center Institutional Animal Care and Use Committee. Mice were housed at the Atlanta VA Medical Center with controlled conditions with free access to food and water. More information on these mice can be found in Appendix.

### Micro-computed tomography scanning, reconstruction, and landmark placement

Ethanol-fixed heads from postnatal day 10 (P10) control (n=6), GOF (n=8), and LOF (n=6) mice and scanned using a SkyScan 1275 micro-computed tomography (micro-CT) system (Bruker, Billerica, Massachusetts, USA). The same parameters were used for each sample. Landmarks were positioned on the skulls and mandibles as previously published (Bamaga et al. 2017; Hassan et al. 2020; Vora et al. 2016). All landmarks were placed by the same blinded observer and intraobserver analysis was conducted to ensure replicability, see the Statistics section. More information on micro-CT scanning, reconstruction, and landmark placement can be found in Appendix.

### Traditional morphometrics using all landmarks

Traditional morphometric analysis was conducted as previously described (Bumann et al. 2022). All results including significant and non-significant measurements are in Appendix. All measurements were compared between groups as true measurements and adjusted for cranial or mandibular centroid size as previously described (Bumann et al. 2022). For more information on the traditional morphometric analysis, including acquiring BMD and bone microarchitecture parameters, see Appendix.

### Assessment of tooth eruption using micro-CT scans

Images of the occlusal surface including bone with teeth and teeth alone were captured using the micro-CT scans of each individual’s left and right mandibular and maxillary molars. Using the images of the teeth alone, each tooth cusp tip, or apex, was identified. Using the image of the bone with teeth, fully visible cusp tips (with no obstructing bone) were manually counted. Fully visible cusp tips were counted by the same observer twice and intraobserver analysis was conducted to ensure replicability, see Appendix.

### Geometric morphometric analyses

Shape differences in the coronoid and condylar processes of all N=20 mandibles were analyzed using geometric morphometrics. For more information on the geometric morphometric analysis, see Appendix.

### TUNEL labeling

TUNEL labeling using the Click-iT™ Plus TUNEL Assay Alexa Fluor 647 (Invitrogen, C10619) was conducted using the manufacturer’s protocol. For more information on tissue sectioning and TUNEL labeling, see Appendix.

### Statistics

For the traditional morphometrics, groups were compared using an unpaired, parametric, two-tailed t test with Welch correction with the False Discovery Rate FDR (Q) set to 5. A stringent p-value of <0.01 was used. For all other t tests, an unpaired, parametric, two-tailed t test and a typical p-value of <0.05 was used. A two-way ANOVA was used to compared TUNEL:DAPI ratio at the three mandibular loci between genotypes. Cohen’s d effect size was used to describe magnitude of difference between groups. All measurements are reported using scatter plots with bars representing mean and standard deviation. See Appendix for more information on statistics.

## RESULTS

*Wnt5a* LOF skull length relative to centroid size was a significant 1.7% shorter than controls (Fig 1D-F, Appendix Figs 1&6B). No body weight differences were detected, indicating differences in body size are not the source of skull size changes (Appendix Fig 2). The trend towards shorter total skull length in *Wnt5a* LOF mice was due in part to midface hypoplasia (Fig 2A). Indicative of midface hypoplasia, *Wnt5a* LOF mice compared to controls had a significant 4.9% decrease in projected upper jaw length (Fig 2A.I) and a significant 10.4% decrease in projected maxilla length (Fig 2A.II). In the transverse dimension, *Wnt5a* LOF mice had a significant 8.3% larger maxillary intermolar width (Fig 2A.III) and a significant 14.5% wider rostral basisphenoid width than controls (Fig 2A.IV). No difference in midface length or transverse cranial widths between *Wnt5a* GOF mice and controls was detected. Interorbital width trended 5.6% wider in *Wnt5a* LOF mice (Appendix Fig 3). Indicative of significant delayed molar eruption, *Wnt5a* LOF mice had 46.5% fewer mandibular (Fig 2B-D) and 36.5% fewer maxillary (Fig 2E-G) molar cusp tips fully unobstructed by alveolar bone at postnatal day 10 than controls (Fig 2H). Though molar morphology changes were not observed (Appendix Fig 4), *Wnt5a* LOF mice had a 6.5% decrease in mandibular molar field length and a 7.7% decrease in maxillary molar field length compared to controls indicating smaller teeth (Appendix Fig 5).

**Figure 2.**
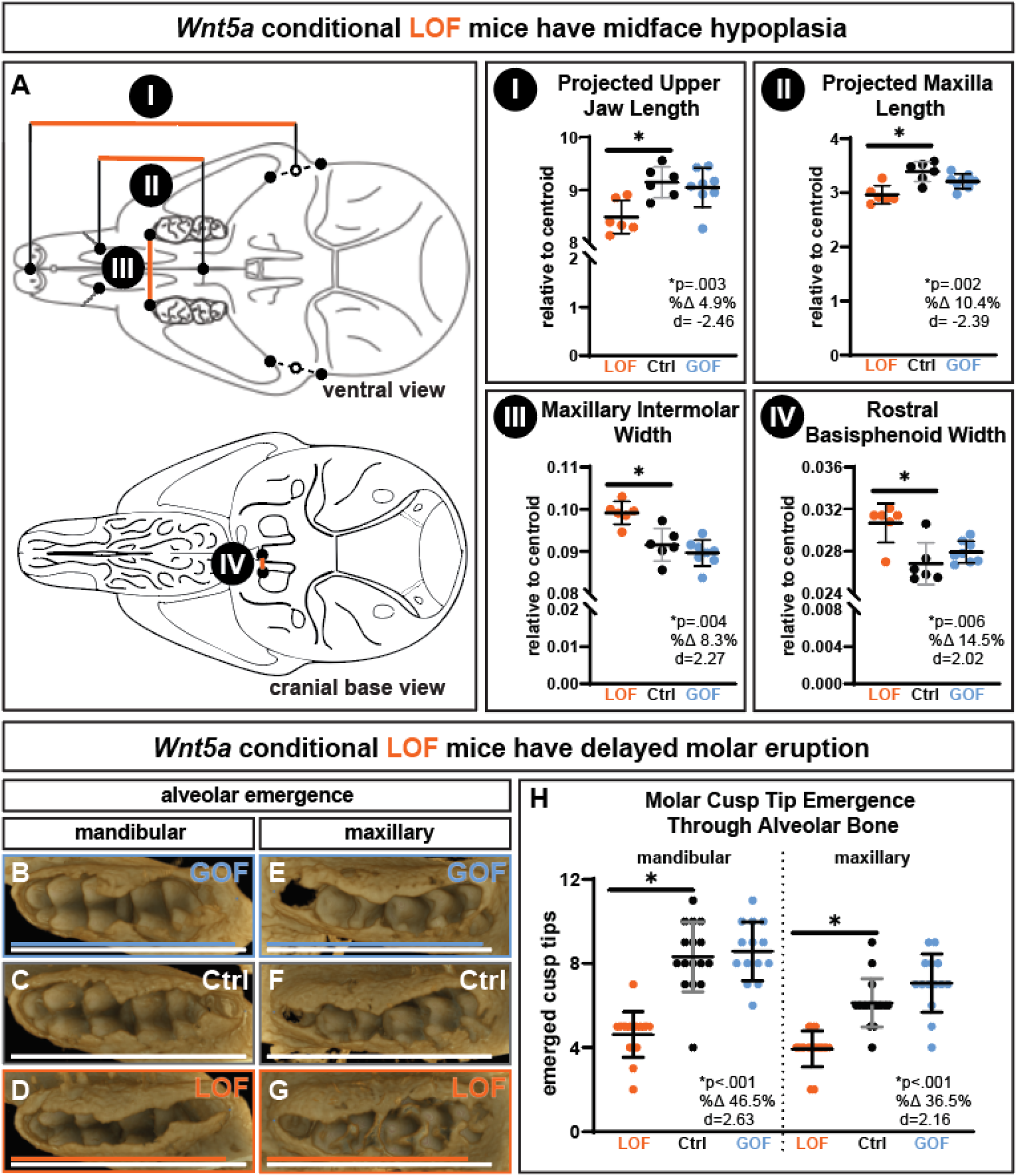
Conditional *Wnt5a* LOF mice have midface hypoplasia and delayed molar eruption. Determined by using anatomical landmarks on micro-CT scans, *Wnt5a* LOF mice alone had a significant **A.I)** decrease in projected upper jaw length, **A.II)** decrease in projected maxilla length, **A.III)** increase in maxillary intermolar width, and **A.IV)** increase in rostral basisphenoid width in the cranial base. The occlusal view of the micro-CT scans demonstrate that in both **B-D)** the mandibular molars and **E-G)** the maxillary molars, bone covers or partially covers more cusp tips in P10 *Wnt5a* LOF mice. Solid lines on the occlusal view micro-CT scans indicate molar field length, which were decreased in the mandible and maxilla in *Wnt5a* LOF mice. **H)** *Wnt5a* LOF mice have fewer mandibular and maxillary molar cusp tips completely uncovered by bone. Exact p-values, percent change, and Cohen’s d effect size are listed within each graph.

Both *Wnt5a* GOF and LOF genotypes had a partially penetrant snout deviation (Fig 3A). Half of the *Wnt5a* GOF mice had a snout deviation: 1/6 had mild asymmetry and 2/6 had a pronounced asymmetry (Fig 3A.V & Fig 3B-C). Slightly less than half of the *Wnt5a* LOF mice had a snout deviation: 1/8 had mild asymmetry and 2/8 had a pronounced asymmetry (Fig 3A.V & Fig 3C-D). All snout deviations in the asymmetrical Wnt5a GOF and LOF were towards the right. There were no observed overt changes in any cranial sutures (data not shown).

**Figure 3.**
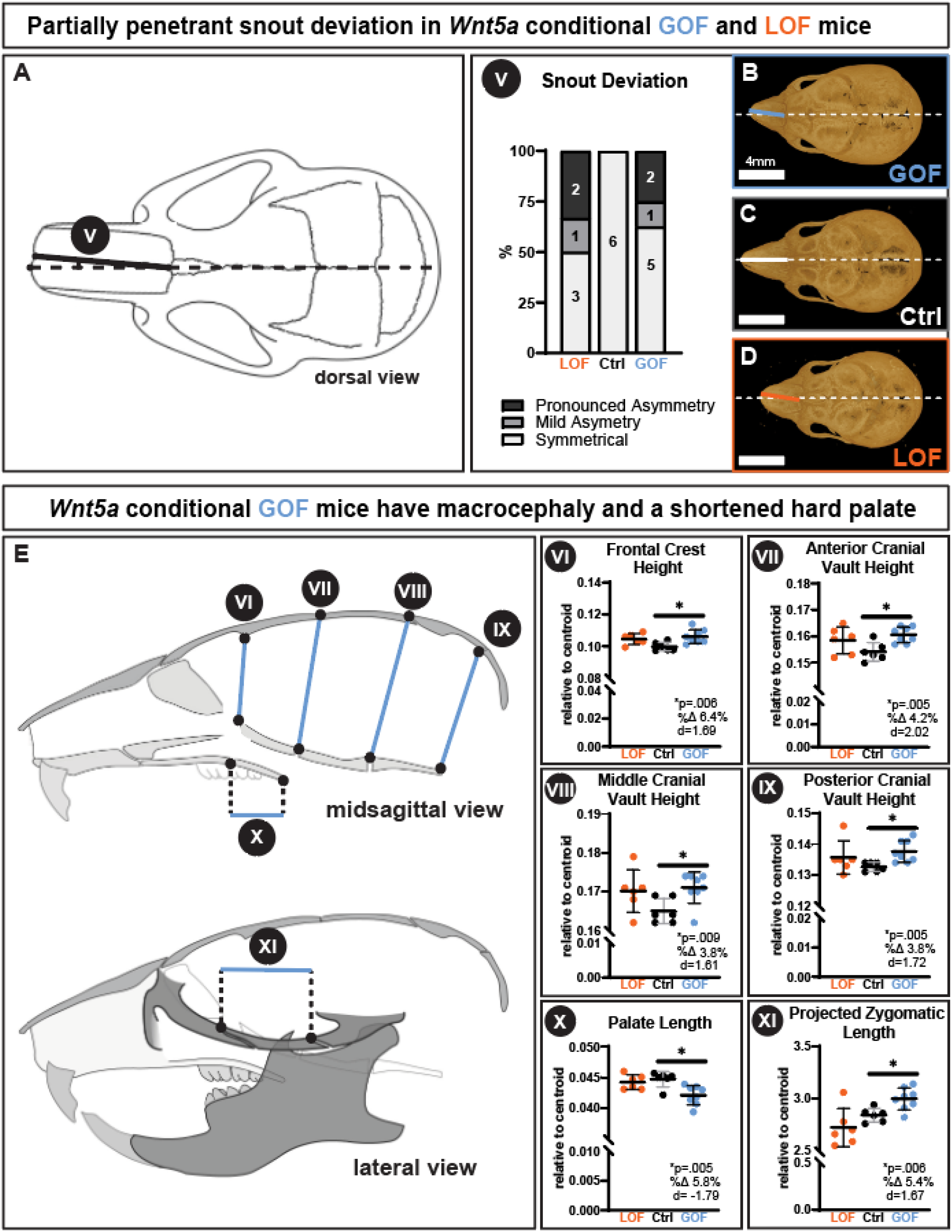
Both conditional *Wnt5a* LOF and GOF mice had a partially penetrant snout deviation, while GOF mice alone had macrocephaly and a shortened hard palate. **A)** Approximately half of the *Wnt5a* LOF and GOF mice had a mild or pronounced snout deviation, indicative of facial asymmetry. **B-D)** Dorsal view micro-CT images demonstrate the snout asymmetry, which deviated to the right in all affected individuals. *Wnt5a* GOF mice alone had a significant **E.VI)** increase in frontal crest height, **E.VII)** increase in anterior cranial vault height, **E.VIII)** increase in middle cranial vault height, **E.IX)** increase in posterior cranial vault height, which taken together indicate macrocephaly. *Wnt5a* GOF mice alone also had a significant **E.X)** decrease in palate length and **E.XI)** increase in projected zygomatic length. Exact p-values, percent change, and Cohen’s d effect size are listed within each graph.

While the *Wnt5a* LOF mice had midface hypoplasia and delayed tooth eruption, *Wnt5a* GOF mice had macrocephaly, shortened hard palate, micrognathia, and mandibular process morphology changes. Indicative of macrocephaly, *Wnt5a* GOF mice had a significant 6.4% larger frontal crest height (Fig 3.VI), a significant 4.2% larger anterior cranial vault height (Fig 5.VII), a significant 3.8% larger middle cranial vault height (Fig 5.VIII), and a significant 3.8% larger posterior cranial vault height (Fig 5.IX) compared to controls. In the anterior-posterior dimension, *Wnt5a* GOF mice had a significant 5.8% shorter palate length than controls (Fig 5.X). However, *Wnt5a* GOF mice had a significant 5.4% longer projected zygomatic length than controls (Fig 5.XI).

Indicative of micrognathia, *Wnt5a* GOF mice had a significant 3.5% decrease in superior mandible length (Fig 4A.XII) and a significant 4.4% decrease in inferior mandible length compared to controls (Fig 4A.XIII). There were no changes in cranial or mandibular BMD (Appendix Fig 6C&D) and centroid (Appendix Fig 6A&B). The only bone microarchitecture change detected in the mandibles was a significant 11% decrease in bone surface and a significant 32.9% decrease in trabecular connectivity in *Wnt5a* GOF mandibles (Appendix Fig 6E&F).

**Figure 4.**
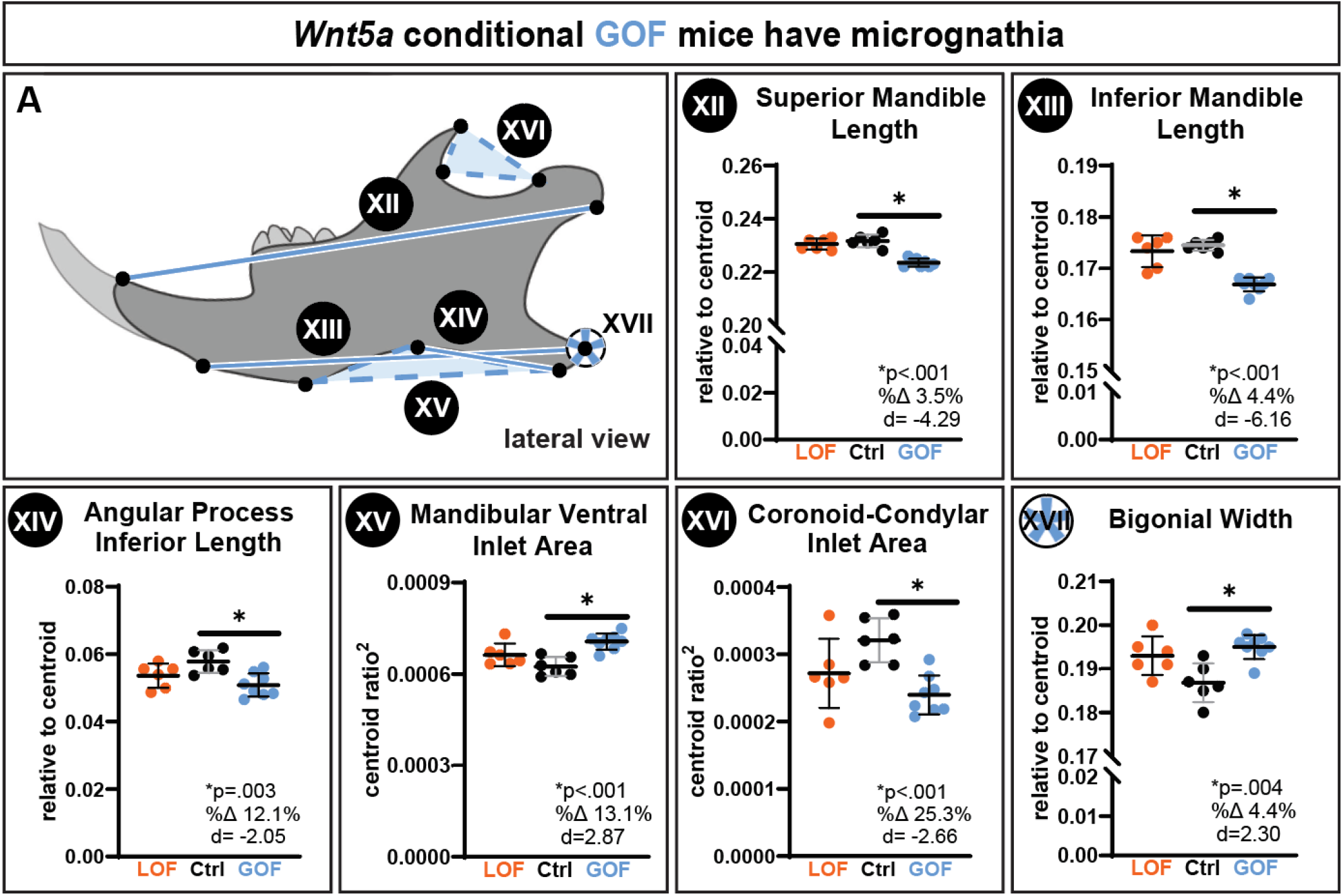
Conditional *Wnt5a* GOF mice had micrognathia, indicated by a significant decrease in **A.XII)** superior mandible length, **A.XIII)** inferior mandible length, and **A.XIV)** angular process inferior length. *Wnt5a* GOF mice also had changes in the mandibular processes, indicated by a significant **A.XV)** increase in mandibular ventral inlet area and **A.XVI)** decrease in coronoid-condylar inlet area. *Wnt5a* GOF mice had **A.XVII)** significantly wider bigonial width between the angular process of the left and right hemimandibles. Exact p-values, percent change, and Cohen’s d effect size are listed within each graph.

As determined by traditional morphometric analysis, *Wnt5a* GOF mice had a significant 12.1% smaller angular process inferior length (Fig 4A.XIV), a significant 13.1% larger mandibular ventral inlet area (Fig 4A.XV), and a significant 25.3% smaller mandibular coronoid-condylar inlet area than controls (Fig 4A.XVI). The width between the left and right mandibular angular process (bigonial width) was significantly 4.4% wider in *Wnt5a* GOF mice (Fig 4A.XVII). Notably, the *Wnt5a* GOF mice did not have a wider cranial base, indicating that the larger bigonial width in the *Wnt5a* GOF mice was not due to a widening of the cranial base.

As determined by geometric morphometric analysis, there were morphological differences in the *Wnt5a* GOF coronoid and condylar processes (Fig 5A-C). Principal component analysis of GPA coordinates indicated principal components 1-4 each captured at least 10% of variation between samples. In *Wnt5a* GOF mice, coronoid processes tended to be shorter and less rounded, condylar processes tended to have a pinched ramus and a rounded articular surface, and the angular process tilted downward (Fig 5D). The full list of principal components are listed in Appendix Table 1, and the morphology captured by principal components 1-4 are visualized in Appendix Table 2. *Wnt5a* GOF mandibular process morphology differed significantly from both controls and *Wnt5a* LOF mice, and there was a significant allometric relationship between mandible centroid size and mandibular process shape in *Wnt5a* GOF mice only (Appendix Table 3). The allometric slope between predicted shape and log mandible centroid size angles sharply downward in *Wnt5a* GOF mice in contrast to the other two genotypes, indicating that smaller *Wnt5a* GOF mandible centroid size correlated with a more drastic mandibular process shape difference from controls (Fig 5E).

**Figure 5.**
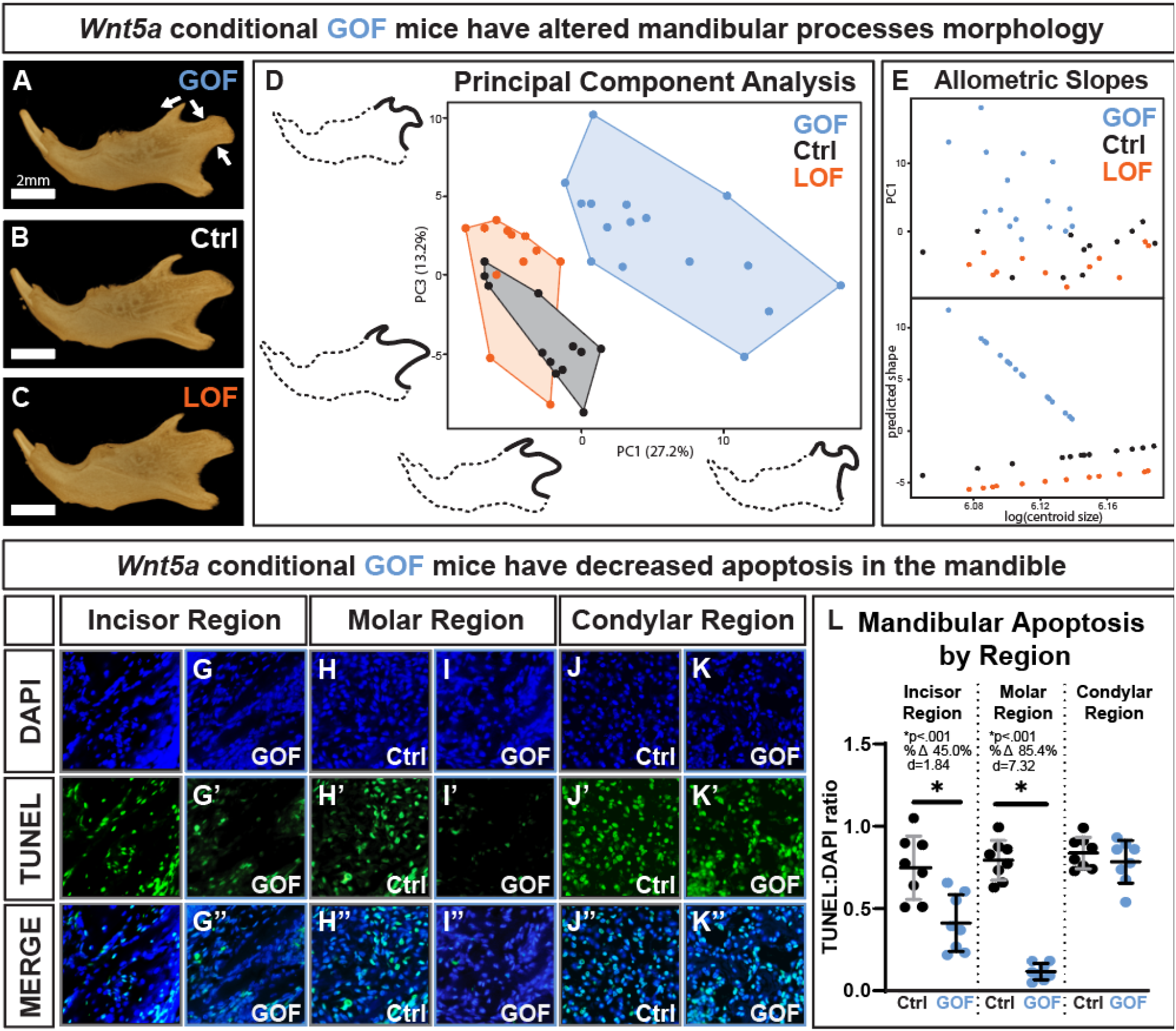
Conditional *Wnt5a* GOF mice had altered morphology of the mandibular processes as determined by geometric morphometrics but have decreased mandibular apoptosis. **A-C)** Lateral view micro-CT images of segmented mandibles demonstrate the shorter and less rounded coronoid process, pinched ramus and rounded articular surface of the condylar process, and the downward tilted angular process in *Wnt5a* GOF mice. **D)** PC1 (describes 27.2% of variation between samples) vs PC3 (describes 13.2% of variation between samples) of the principal component analysis of the mandibular process geometric morphometrics; outlines demonstrating phenotype are 2x exaggerated and to scale. **E)** There is an allometric relationship between mandible size and mandibular process shape in the *Wnt5a* GOF mice alone. **F-K)** Representative images of control and Wnt5a GOF mice stained for DAPI (blue) and TUNEL-indicated apoptosis (green). **L)** The TUNEL:DAPI ratio was significantly decreased in *Wnt5a* GOF mice in mandibular tissue near the incisor and near the molars but was similar to controls in the mandibular condyle.

Indicated by TUNEL staining (Fig 5F-K), *Wnt5a* GOF mice had a significant 45% decrease in apoptosis in mandibular bone near the incisor and an 85% decrease in apoptosis in mandibular bone near the molars compared to controls (Fig 5L). *Wnt5a* GOF mice had similar levels of TUNEL staining to controls in the mandibular condylar process.

## DISCUSSION

One key finding of this study was that conditional LOF and conditional GOF of *Wnt5a* result in distinct craniofacial phenotypes reminiscent of the heterogeneity of craniofacial phenotypes in individuals with Robinow syndrome. Craniofacial anomalies noted in patients with Robinow Syndrome that were identified in this study included midface hypoplasia and a trend toward hypertelorism only in the *Wnt5a* LOF group, while macrocephaly, shortened hard palate, and micrognathia were noted only in the *Wnt5a* GOF group (key findings are summarized in Appendix Table 6). These craniofacial anomalies are all associated with Robinow syndrome, but do not necessarily all co-occur in the same individuals (Conlon et al. 2021). Our finding of a function-specific effect of *Wnt5a* is supported by other *Wnt5a* mouse models. For example, conditional *Wnt5a* LOF but not GOF in *Cdx2*^*cre*^, which targets posterior body structures, mice results in severe hindlimb and tail truncation (Simonson et al. 2022).

Another key finding of this study was the identification of several craniofacial and dental phenotypes not currently clinically associated with *WNT5A* mutations: tooth eruption delay, TMJ changes, and facial asymmetry. Delayed molar eruption was noted only in the *Wnt5a* LOF group, while mandibular process morphology changes were noted only in the *Wnt5a* GOF group. Temporomandibular joint disorders in individuals with Robinow syndrome are not reported in the literature, though there is an association between *Wnt5a* signaling and TMJ dysfunction in other animal models (Yang et al. 2015). Facial asymmetries were detected in both the *Wnt5a* LOF and GOF mice groups. Craniofacial asymmetries have not been reported in individuals with *WNT5A*-associated Robinow syndrome, but clinical photographs of individuals with Robinow syndrome suggest asymmetries may be present but not documented (Conlon et al. 2021). WNT5A is involved in planar cell polarity and is needed for proper left-right patterning in early embryonic development, which may explain why some *Wnt5a* LOF and GOF mice displayed asymmetries in the midface and should be explored further (Hosseini-Farahabadi et al. 2017; Minegishi et al. 2017). Delayed molar eruption, mandibular process morphology, and craniofacial asymmetries should be assessed in patients with Robinow Syndrome.

Our findings are consistent with previous reports that *Wnt5a* GOF results in a shorter lower jaw (Hosseini-Farahabadi et al. 2017). Viral overexpression of wild type *WNT5A* and *WNT5A* with a missense variant in chick lower jaw results in a 20% reduction in lower jaw length (Hosseini-Farahabadi et al. 2017). Conditional *Wnt5a* GOF, viral *Wnt5a* overexpression (Hosseini-Farahabadi et al. 2017), viral missense variant-containing *Wnt5a* overexpression (Hosseini-Farahabadi et al. 2017), and global *Wnt5a* knockout (Yamaguchi et al. 1999) cause micrognathia, but conditional *Wnt5a* LOF does not. This suggests *Wnt5a* LOF may be inconsequential in *Ctsk*-expressing periosteal cells, osteoclasts, pericytes, bone marrow stromal cells, and fibrocartilage in establishing mandibular size and shape, but essential in other cell types during jaw development. We found GOF of *Wnt5a* in the aforementioned cell types disrupts critical processes required for jaw formation. Consequently, mandibular manifestations of *WNT5A* mutations are variant consequence-dependent consistent with micrognathia occurring in only 33-57% of individuals with Robinow syndrome (Conlon et al. 2021; Person et al. 2010).

Cleft lip with/without cleft palate occurs in approximately 40-60% of individuals with Robinow syndrome (Conlon et al. 2021; White et al. 2018; Zhang et al. 2022), but we did not detect clefting in this animal model nor did we detect a difference in perinatal lethality between genotypes. However, conditional *Wnt5a* LOF mice had increased transverse maxillary intermolar width as is observed in unaffected family members of individuals with cleft lip/palate (Weinberg et al. 2006). It is possible that conditional *Wnt5a* LOF may result in heightened vulnerability to cleft palate.

Global Wnt5a knockout (Lin et al. 2011) and *Ctsk*^cre^;*DTA*^*fl/+*^ (Hassan et al. 2022) mice have altered molar morphology, but our conditional *Wnt5a* LOF and GOF mice did not. This suggests *Wnt5a* affects tooth morphology, but not in *Ctsk* lineage cells. Conversely, *Ctsk* lineage cells affect tooth morphology via *Wnt5a-*independent mechanisms. Though we did not observe tooth morphology changes, we detected delayed molar eruption in conditional *Wnt5a* LOF mice. *Wnt5a* is known to be robustly expressed in alveolar bone during tooth eruption and promote osteoclastogenesis (Maeda et al. 2012; Xiang et al. 2014), and our data provide a direct link between bone *Wnt5a* ablation and delayed tooth eruption. Additionally, *Wnt5a* LOF mice had decreased mandibular and maxillary molar field length, however this magnitude of decreased molar size would not meet the clinical criteria for microdontia.

Interestingly, the phenotype of our conditional *Wnt5a* GOF mice closely resembled that of *Ctsk*^cre^;*DTA*^*fl/+*^ mice, with both exhibiting shortened mandibles, altered mandibular condyles, and enlarged cranial vaults (Hassan et al. 2022). We consequently hypothesized that excess *Wnt5a* results in cell death in affected cells, explaining why loss of *Ctsk*^*cre*^-expressing cells and *Wnt5a* GOF in *Ctsk*^*cre*^-expressing cells results in a similar craniofacial phenotype. Unexpectedly, *Wnt5a* GOF mice had significantly decreased cell death in mandibular tissue. *Wnt5a* has a well-established association with decreased cell death, including in craniofacial tissue (Bo et al. 2016; Vuga et al. 2009; Zhu et al. 2025). It is possible physiological cell death during jaw development was disrupted in *Wnt5a* GOF mice resulting in craniofacial changes. Additionally, the *Ctsk*^cre^;*DTA*^*fl/+*^ study and our own investigated slightly different cell populations, as the *Ctsk*^cre^;*DTA*^*fl/+*^ study utilized a different *Ctsk*^cre^ mouseline (Chiu et al. 2004). Further investigation using additional techniques to assess cell death is needed to definitively establish a connection between these phenomena.

Previously published data show male *Wnt5a* LOF mice have 38% lower femur BMD than controls (Roberts et al. 2020), but interestingly, we detected no difference in mandible BMD between groups (Appendix Fig 6C). This may explain why some individuals with *WNT5A*-associated Robinow syndrome have mild osteopenia of the non-cranial skeleton but normal BMD in the cranial skeleton (Shayota et al. 2020). Bone surface and connectivity, which were significantly different in *Wnt5a* GOF mandibles, did not differ from controls in *Wnt5a* LOF mandibles in agreement with no reported difference in these measures in *Wnt5a* LOF femurs (Roberts et al. 2020). However, other bone microarchitecture parameters were altered in the femurs of male *Wnt5a* LOF mice but did not differ between genotypes in the mandible (Roberts et al. 2020). Similarly, *Wnt5a*^*fl/fl*^*;Osx*^*cre*^ mice have decreased trabecular separation in the femur, but data in the mandible in this model is not available (Maeda et al. 2019). The contrasting BMD and bone microarchitecture phenotypes between *Wnt5a* LOF femurs and mandibles indicate a bone-specific effect of *Wnt5a* loss-of-function in *Ctsk*^*cre*^-expressing cells.

In conclusion, the craniofacial and dental phenotype differed between mice with conditional GOF and LOF of *Wnt5a* in *Ctsk*^*cre*^-expressing cells, consistent with other preclinical reports and the craniofacial phenotype heterogeneity in *Wnt5a*-associated Robinow syndrome. Further, we detected tooth eruption delay, TMJ changes, and facial asymmetry in mice with altered *Wnt5a* that have not been previously reported but for which patients with Robinow syndrome should be monitored. Lastly, *Wnt5a* may play a bone-specific role as demonstrated by differences in mandibles and femurs of conditional *Wnt5a* knockout and in patients with Robinow Syndrome.

## ACKNOWLEDGEMENTS

We thank S.R. Vora for sharing the mouse skull outlines, G. Liu for assistance with animal breeding, B.A. Smith for assistance with data collection, M. Hassan for discussions about geometric morphometric analysis, J.M. Scott for discussions around statistical analysis, and S.L. Dallas, Y. Xie, D. Moore, T.C. Cox, and R. Anderson for helpful conversations.

