## Appendix for "Wnt5a gain- and loss-of-function present distinctly in craniofacial bone"

**Appendix Contents**

Supplemental Materials and Methods.......2-5
Supplemental Figures..............................6-12
Supplemental Tables..............................13-20

### Supplemental Materials and Methods

Animals

*Ctsk*$^{cre}$ mice were acquired from the University of Tokyo (Nakamura et al. 2007) and have been backcrossed into a C57BL/6J background. *Wnt5a*$^{fl/fl}$ (Strain #: 026626) mice were purchased from Jackson Laboratory (Bar Harbor, ME). Conditional *Wnt5a* knockout (LOF) mice were generated as previously reported (Fig 1B&C) (Roberts et al. 2020). *Rosa26-LSL-Wnt5a* mice were acquired from Washington University where they were originally generated (Chen et al. 2014). Homozygous *Rosa26-LSL-Wnt5a* mice were crossed with heterozygous *Ctsk*$^{cre}$ to generate condition *Wnt5a* knock-in (GOF) mice (Fig 1A). *Wnt5a* LOF and *Wnt5a* GOF mice were compared to *Wnt5a*$^{fl/fl}$ controls. All three genotypes are on a C57BL/6J background. Six or more mice per genotype were analyzed to generate a sufficient sample size for craniofacial morphometrics while minimizing excess animal use. All described methods were conducted on mixed sex groups containing males and females. Sex differences were not analyzed as the sample size per genotype per sex did not sufficiently power statistical comparison between groups stratified by sex. The only inclusion criteria was life at postnatal day 10; no difference in pup viability was noted between groups.

Mice were maintained in accordance with applicable state and federal guidelines and all experimental procedures were approved by the Atlanta VA Medical Center Institutional Animal Care and Use Committee (Protocol number: V008-23). Mice were housed at the Atlanta VA Medical Center with controlled conditions (21-24°C temperature; 40-70% humidity; 12h/12h light/dark cycle) with free access to Teklad #2018S food and water (0.1µm filtered).

Micro-computed tomography scanning, reconstruction, and landmark placement

Ethanol-fixed heads from postnatal day 10 (P10) control (n=6), GOF (n=8), and LOF (n=6) mice and scanned using a SkyScan 1275 micro-computed tomography (micro-CT) system (Bruker, Billerica, Massachusetts, USA). A custom 0.5mm aluminum filter was used. Samples were scanned at a voltage of 55kV and a current of 180µA with an integration time of 45ms and a resolution of 14µm$^3$. Samples were scanned in 180° at a rotation step of 0.3°.

Reconstruction was completed using NRecon software version 1.7.4.2 (Bruker, Billerica, Massachusetts, USA). Gaussian smoothing, ring-artifact reduction, and beam-hardening correction were applied, and the same thresholding parameters were used for each sample. Three-dimensional rendering and landmark placement were performed using Drishti software version 2.6.5 (National Computational Infrastructure's VizLab, Canberra, Australia). Landmarks were positioned on the skulls and mandibles as previously published (Bamaga et al. 2017; Hassan et al. 2020; Vora et al. 2016). All landmarks were placed by the same blinded observer and intraobserver analysis was conducted to ensure replicability, see the Statistics section.

Traditional morphometrics using all landmarks

Euclidean measurements, including angles and linear distances, between landmark points were taken for all measurements. Projected measurements were found by calculating the distance between parallel planes intersecting landmark points, as previously described (Vora et al. 2016). The mean of right and left measurements was used to compare all bilaterally paired landmarks between groups. All results including significant and non-significant measurements are in Appendix Table 4. All measurements were compared between groups as true measurements and adjusted for cranial or mandibular centroid size in order to mitigate any differences in skull size. Centroid size of the crania and mandibles were calculated using the root centroid size equation, $RCS = \sqrt{\sum_{i=1}^{n}(x_i - \bar{x})^2 + (y_; - \bar{y})^2 + (z_i - \bar{z})^2}$ (MacLeod 2008). Each measurement is then a ratio of the size of the cranium or mandible, and comparisons can be made between groups (Klingenberg 2016; Maga et al. 2015).

Snout asymmetry was analyzed by measuring the angle of deviation between the mid-sagittal plane and the internasal suture. Landmarks were placed at the anterior point of the nasal bone intersecting with the mid-sagittal plane, the nasion, and the anterior point of the nasal bone. An angular measurement was taken with the nasion as the vertex. Mild asymmetry was assigned if the value fell outside the range of asymmetry in the control group but was less than twice the most asymmetrical control. Pronounced asymmetry was assigned if the value was greater than twice the most asymmetrical control.

<u>BMD and bone microarchitecture</u>

Bone mineral density (BMD) and microarchitecture parameters were determined for all N=20 samples using CT Analyser software version 1.17.7.2 + (Bruker). Calibration was done using Skyscan 0.30 and 1.25 g/cm3 CaHA mouse-sized (2mm diameter, 10mm length) density phantoms (Bruker). Measurements were analyzed for a Region of Interest (ROI) including the full skull, and with the ROI selected for the mandible only. A density threshold of 40-255 Hounsfield units (HU) was applied to the full skull ROI, and a density threshold of 25-255 HU was applied to the mandible ROI. The ROI was shrink-wrapped, and black and white speckles of greater than 10 voxels were removed.

<u>Geometric morphometric analyses</u>

Shape differences in the coronoid and condylar processes of all N=20 mandibles were analyzed using geometric morphometrics. The left and right side of each mandible were analyzed in case asymmetries were noted. Lateral images of the right and left mandible were captured from the 3D reconstructions of the skull micro-CT scans, including the location of the 11 landmarks placed on the mandible in 3D as described in the geometric morphometrics analyses section of the main methods, which were used to estimate mandible centroid size. Images of the right side of the mandible were mirrored to allow direct mandibular process shape comparison of both sides. A blinded observer placed 59 semi-landmarks on each image using tpsDig2 (v 2.32) (Rohlf 2006). Two landmarks were also placed on either side of a scale bar for quality control to ensure all landmarks were scaled equally. The full description of mandibular 3D-placed landmarks and semi-landmarks placement are in Appendix Fig 7.

Landmark coordinates were analyzed using the *Geomorph* R package (Adams and Otárola-Castillo 2013). Generalized Procrustes analysis (GPA) using the *gpagen* function was used to align the landmark coordinates to adjust for differences in scale, orientation, and position.

Procrustes coordinates from a GPA using all 3D-placed and semi-landmarks were used to assess shape variation using a Principal Component Analysis (PCA) with the *gm.prcomp* function. Only the principal component axes that represented greater than ten percent of the shape variation were used in the multivariate analysis of covariance (MANCOVA) models; using *lm.rrpp* function (type III sums of squares, 10,000 iterations). Genotype, sex, side (right/left), and individual, and all their interactions were used as categorical independent variables, and log-transformed mandible centroid size as the covariate. Variables that were not significant were removed from the final model. The final model included only genotype and mandible centroid size. A pairwise test was used to look at the effect of genotype on shape using the *pairwise* function, which measured the Euclidean distance between group means (10,000 iterations, 0.95 confidence). The interactions between log transformed mandible centroid size and each genotype were used to test for allometric relationships with shape. A pairwise test using the *pairwise* function measured the distance between slope vectors for each genotype to determine if any genotype had an allometric slope that significantly differed from the others (10,000 iterations, 0.95 confidence).

Tissue sectioning and TUNEL labeling

Formalin-fixed whole heads were decalcified in 10% EDTA (pH 7.2) for 10 days. Tissues were then embedded in paraffin blocks and were stored at 4°C. Blocks were placed in a beaker and chilled in an ice box to maintain low temperature during sectioning, which prevented tissue breakage. Using a microtome, control and experimental mice head tissues were sectioned transversely and sagittally. Serial sections of mice head tissues were cut in the horizontal plane at a thickness 7 μm and placed on a distilled water bath at 42°C. Mice head sections were transferred from distilled water bath to superfrost/plus microscope slides for processing and staining. TUNEL labeling using the Click-iT Plus TUNEL Assay Alexa Fluor 647 (Invitrogen, C10619) was conducted using the manufacturer's protocol on 7μm formalin fixed paraffin embedded transverse sections of control and *Wnt5a* GOF mice. Four loci in three anatomical regions were assessed for TUNEL per individual: mandibular bone sans teeth near the mandibular incisor, mandibular bone sans teeth near the mandibular molars including angular process cartilage, and mandibular condyle bone and cartilage (n=2 mice/genotype, 3 sections/mouse, 4 loci/section). TUNEL-stained sections were mounted in DAPI mounting media (Vector Laboratories, H-1200-10), imaged the following day to prevent signal loss on a Keyence BZ-X810 Epifluorescence High Content Imaging System (Keyence, Itasca, IL), and DAPI-positivity and TUNEL-positivity was quantified using FIJI ImageJ (Schindelin et al. 2012).

Statistics

Outcome measures were traditional craniofacial morphometrics (primary outcome), tooth emergence, mandibular BMD and bone microarchitecture, geometric morphometrics of the mandibular processes, and TUNEL-indicated DNA damage. All comparisons except geometric morphometrics were done in GraphPad Prism v10.4.1 (GraphPad, San Diego, CA, USA). For the traditional craniofacial morphometrics, groups were compared using an unpaired, parametric, two-tailed t test with Welch correction to evaluate statistical significance, with the False Discovery Rate FDR (Q) set to 5. *Wnt5a* GOF were compared to control mice and *Wnt5a* LOF were compared to control mice, but *Wnt5a* GOF and *Wnt5a* LOF mice were not compared to one another. A stringent p-value of $<0.01$ was used to determine if the traditional craniofacial morphometrics measurement likely meaningfully differed between groups. For all other t tests, an unpaired, parametric, two-tailed t test and a typical p-value of $<0.05$ was used. A two-way ANOVA was used to compare TUNEL:DAPI ratio between the three mandibular loci and between genotypes and a typical p-value of $<0.05$ was used. All measurements were reported using scatter plots with bars representing mean and standard deviation. Cohen's d effect size was used to describe magnitude of difference between groups. No animals were excluded from the analysis.

Three randomly selected samples from each genotype (N=9) were landmarked a second time by the same observer to ascertain traditional morphometrics reliability. Measurements were considered replicable when the coefficient of variation between the two landmark sets was less than 10%. Appendix Table 5 contains the intraobserver coefficient of variation for all 114 measurements.

To ascertain the reliability of the cusp apex counting methodology, all individuals were assessed twice and the two observations were compared using intraclass correlation coefficients (ICC) in IBM SPSS Statistics (Version 29). The appropriate model for this experimental design is a two-way mixed-effects model, single rater type, absolute agreement definition (Koo and Li 2016). The cusp apex counting methodology ICC 95% confidence interval was .909 to .963, indicating excellent method reliability (Koo and Li 2016).

Supplemental Materials and Methods References

Adams DC, Otárola-Castillo E. 2013. Geomorph: An r package for the collection and analysis of geometric morphometric shape data. Methods in ecology and evolution. 4(4):393-399.

Bamaga I, O'Sullivan R, Schmitz J, Fath W, Fajardo R. 2017. An osteoblast origin for craniofacial dysplasia in neurofibromatosis type i. J Dent Health Oral Disord Ther. 6(6):00223.

Chen J, Tu X, Esen E, Joeng KS, Lin C, Arbeit JM, Rüegg MA, Hall MN, Ma L, Long F. 2014. Wnt7b promotes bone formation in part through mtorc1. PLoS genetics. 10(1):e1004145.

Hassan MG, Kaler H, Zhang B, Cox TC, Young N, Jheon AH. 2020. Effects of multi-generational soft diet consumption on mouse craniofacial morphology. Frontiers in physiology.783.

Klingenberg CP. 2016. Size, shape, and form: Concepts of allometry in geometric morphometrics. Development genes and evolution. 226(3):113-137.

Koo TK, Li MY. 2016. A guideline of selecting and reporting intraclass correlation coefficients for reliability research. Journal of chiropractic medicine. 15(2):155-163.

MacLeod N. 2008. Palaeomath: Part 15-size & shape coordinates. The Palaeontological Association: Palaeontology Newsletter. 69.

Maga AM, Navarro N, Cunningham ML, Cox TC. 2015. Quantitative trait loci affecting the 3d skull shape and size in mouse and prioritization of candidate genes in-silico. Frontiers in Physiology. 6.

Nakamura T, Imai Y, Matsumoto T, Sato S, Takeuchi K, Igarashi K, Harada Y, Azuma Y, Krust A, Yamamoto Y et al. 2007. Estrogen prevents bone loss via estrogen receptor α and induction of fas ligand in osteoclasts. Cell. 130(5):811-823.

Roberts JL, Liu G, Paglia DN, Kinter CW, Fernandes LM, Lorenzo J, Hansen MF, Arif A, Drissi H. 2020. Deletion of wnt5a in osteoclasts results in bone loss through decreased bone formation. Annals of the New York Academy of Sciences. 1463(1):45-59.

Rohlf FJ. 2006. Tpsdig, version 2.10. http://life bio sunysb edu/morph/index html.

Schindelin J, Arganda-Carreras I, Frise E, Kaynig V, Longair M, Pietzsch T, Preibisch S, Rueden C, Saalfeld S, Schmid B et al. 2012. Fiji: An open-source platform for biological-image analysis. Nature Methods. 9(7):676-682.

Vora SR, Camci ED, Cox TC. 2016. Postnatal ontogeny of the cranial base and craniofacial skeleton in male c57bl/6j mice: A reference standard for quantitative analysis. Frontiers in Physiology. 6:417.

### Supplemental Figures

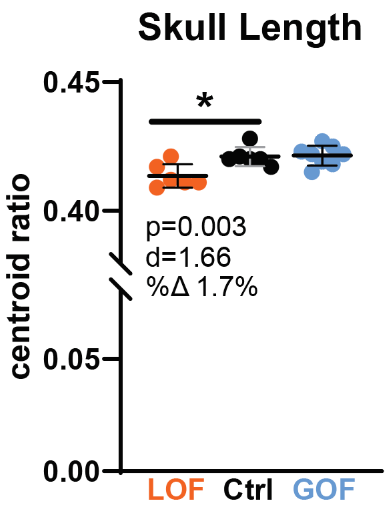

**Appendix Figure 1**. Total skull length was a significant 1.7% smaller in *Wnt5a* LOF mice.

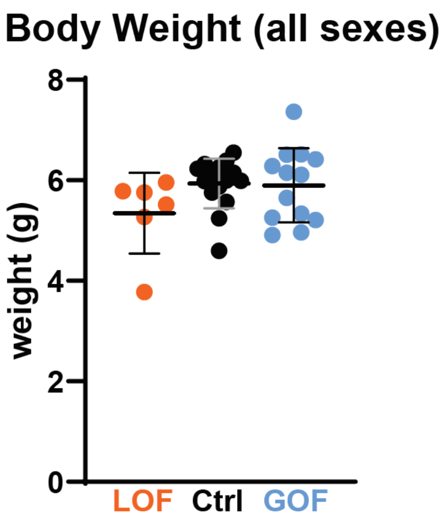

**Appendix Figure 2.** Body weight did not differ between groups.

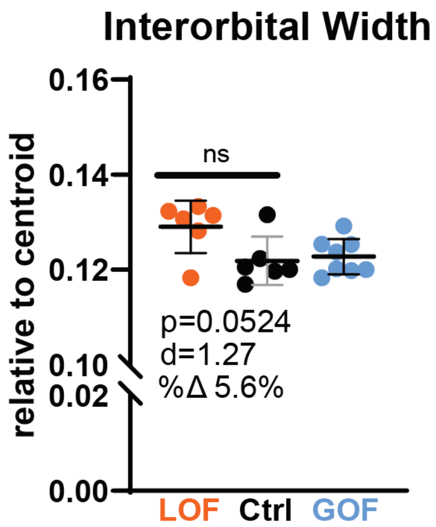

**Appendix Figure 3**. Interorbital width, indicative of hypertelorism, trended up in *Wnt5a* GOF mice but did not meet the threshold of statistical significance using a false discovery rate.

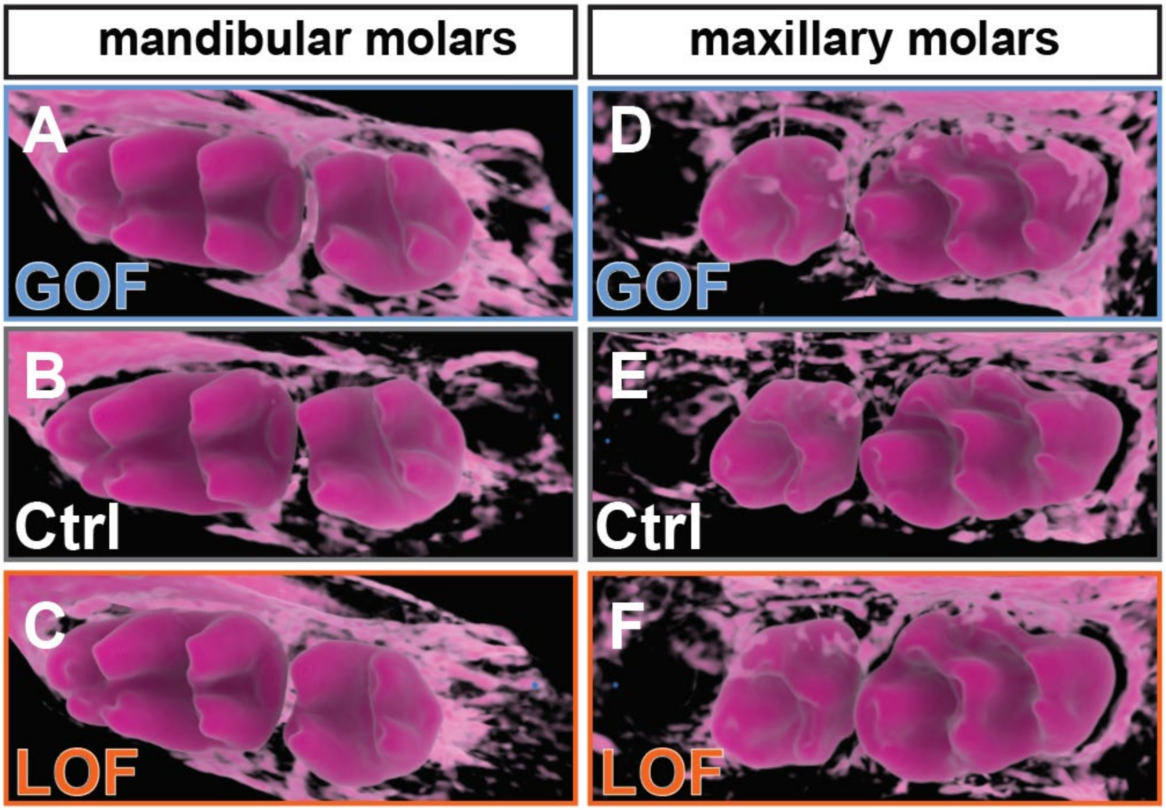

**Appendix Figure 4**. Morphology changes were not observed in the dentition. **A-C)** mandibular and **D-F)** maxillary molars were similar in all genotypes. Images are pseudocolored pink for visualization of morphology, but color does <u>not</u> indicate mineral density.

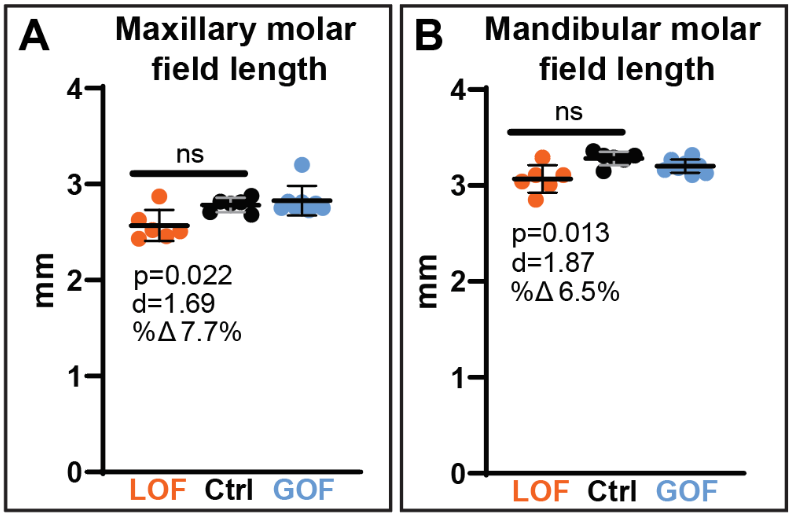

**Appendix Figure 5.** A) Wnt5a LOF mice had a 7.7% shorter maxillary molar field length and B) a 6.5% shorter mandibular molar field length.

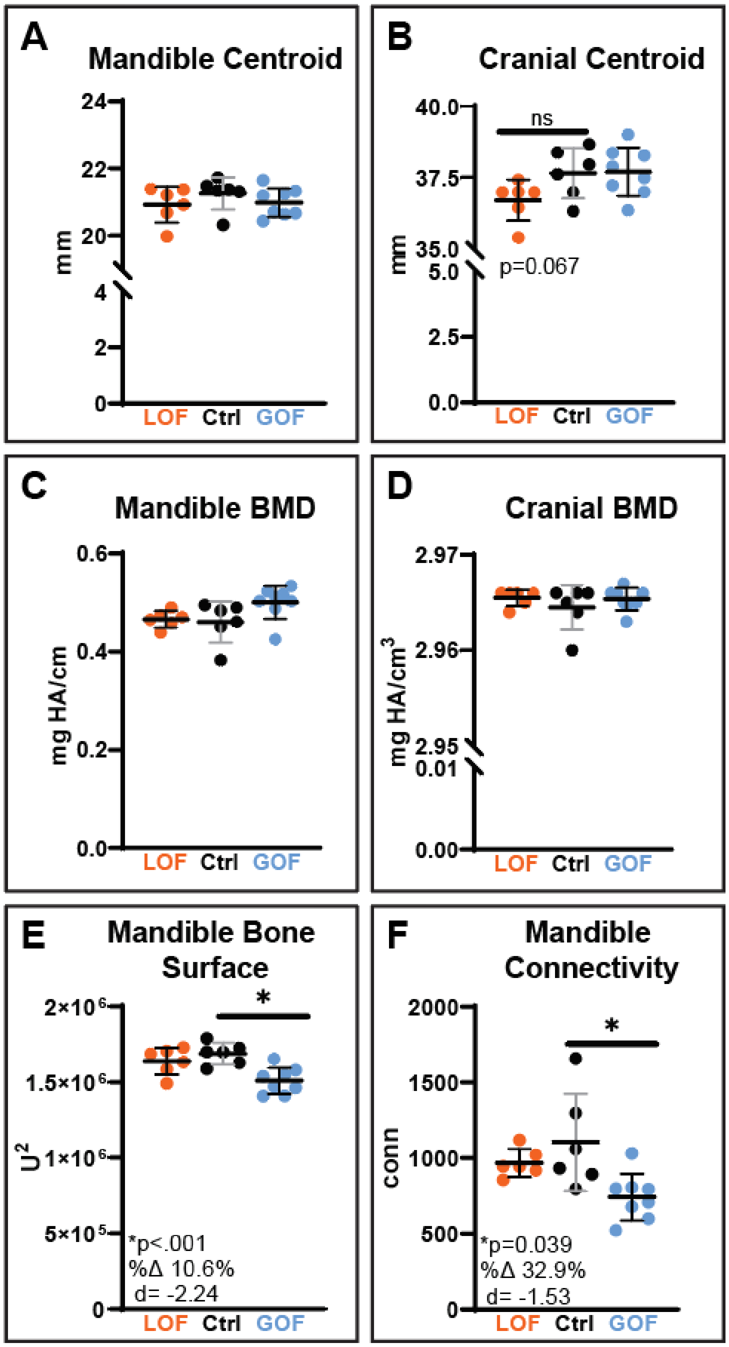

**Appendix Figure 6**. A) Mandible centroid size did not differ between genotypes, but B) cranial centroid size trended down in *Wnt5a* LOF mice. C) Mandible BMD and D) cranial BMD did not differ between groups. E) Mandible bone surface was a significant 10.6% less in *Wnt5a* GOF mice. F) Mandible connectivity was a significant 32.9% less in *Wnt5a* GOF mice.

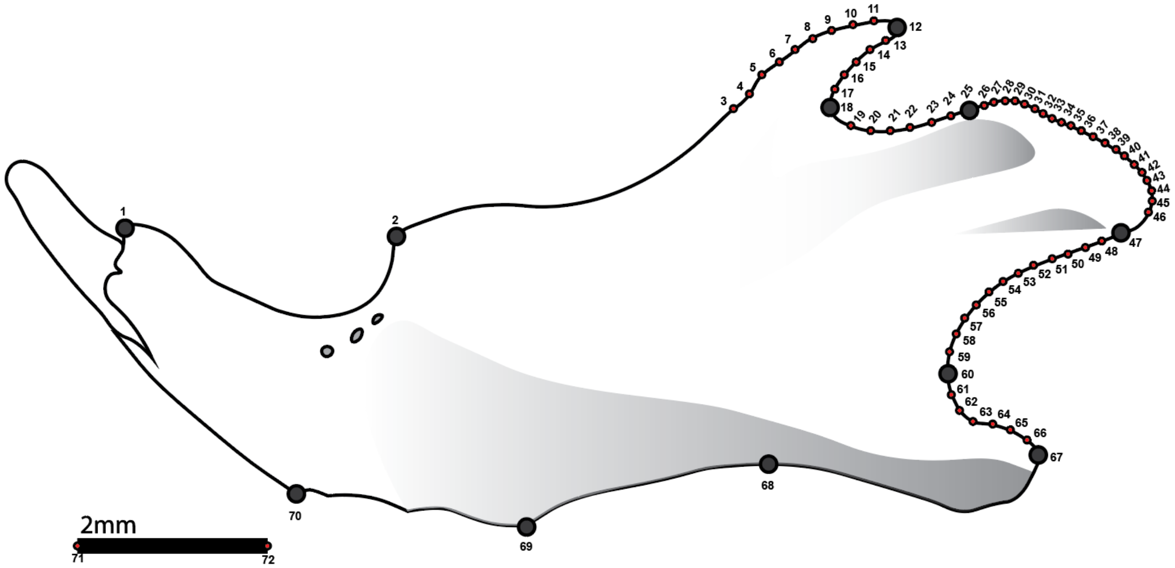

| 3D LM | 2D LM | Type | Landmark Descriptions |
|---|---|---|---|
| M01 | 1 | | Most superior point of incisor alveolus (Midpoint on alveolar bone lingual to the mandibular incisor) |
| M02 | 2 | | Most anterior point of the first molar alveolus |
| | 3-11 | Semi- | A line was drawn between 2(M02) and 12(M04) and divided into three equal sections. Semi-landmarks 3-11 were placed in the 3rd section, on this line along the anterior curve of the coronoid process. |
| M04 | 12 | | Most posterior tip of the coronoid process |
| | 13-17 | Semi- | A line was drawn between 12(M04) and 18(M05). Semi-landmarks 13-17 were placed equidistant on this line along the posterior curve of the coronoid process. |
| M05 | 18 | | Most anterior/inferior concave point of the coronoid process |
| | 19-24 | Semi- | A line was drawn between 18(M05) and 25(M06). Semi-landmarks 19-24 were placed equidistant on this line along the dorsal curve of the neck of the condylar process. |
| M06 | 25 | | Most anterior point of the articular surface of the condyle |
| | 26-46 | Semi- | A line was drawn between 25(M06) and 47(M07) and divided into two equal sections. Landmark 36 was placed on the center of this line, along the curve of the condylar process. A line was drawn between landmarks 25 and 36, and semi-landmarks were placed equidistant on this line, along the curve of the condylar process. This was then repeated for semi-landmarks placed between 36 and 47 along the curve of the condylar process. |
| M07 | 47 | | Most posterior tip of the condyle (Posterior inferior point on mandibular condyle) |
| | 48-59 | Semi- | A line was drawn between 47(M07) and 60(M08). Semi-landmarks 48-59 were placed equidistant on this line along the ventral curve of the neck of the condylar process. |
| M08 | 60 | | Most anterior concave point between the condyle and the angle of mandible |
| | 61-66 | Semi- | A line was drawn between 60(M08) and 67(M09). Semi-landmarks 61-66 were placed equidistant on this line along the dorsal curve of the neck of the angular process. |
| M09 | 67 | | Most posterior tip of the mandibular angle (Posterior tip of the angular process) |
| M11 | 68 | | Ascending ramus dorsal-most ventral point |
| M12 | 69 | | Most inferior point of the alveolar region |
| M13 | 70 | | Anterior inferior most point on the body of the mandible |
| | 71-72 | Scale Bar | Landmarks are placed at the ends of the scale bar to use for resizing landmarks to correct for size variations in screenshots. |

**Appendix Figure 7.** Three-dimensionally placed landmarks (large dots) and two-dimensionally placed semi-landmarks (small dots) used for mandible processes geometric morphometric analysis, and description of all mandibular landmarks placed for geometric morphometric analysis of mandible processes shape. LM=landmark

### Supplemental Tables

| Principal Component | Eigen-values | Proportion of Variance | Cumulative Proportion |
|---|---|---|---|
| **PC1** | 38.134 | 27.238 | 27.238 |
| **PC2** | 24.800 | 17.714 | 44.952 |
| **PC3** | 18.420 | 13.157 | 58.109 |
| **PC4** | 14.020 | 10.014 | 68.124 |
| **PC5** | 6.643 | 4.745 | 72.869 |
| **PC6** | 5.612 | 4.008 | 76.877 |
| **PC7** | 4.603 | 3.288 | 80.165 |
| **PC8** | 4.198 | 2.998 | 83.164 |
| **PC9** | 3.081 | 2.201 | 85.364 |
| **PC10** | 2.346 | 1.675 | 87.040 |
| **PC11** | 2.115 | 1.511 | 88.550 |
| **PC12** | 1.702 | 1.216 | 89.766 |
| **PC13** | 1.549 | 1.106 | 90.872 |
| **PC14** | 1.419 | 1.013 | 91.886 |
| **PC15** | 1.264 | 0.903 | 92.788 |
| **PC16** | 1.083 | 0.773 | 93.562 |
| **PC17** | 0.985 | 0.703 | 94.265 |
| **PC18** | 0.841 | 0.601 | 94.866 |
| **PC19** | 0.819 | 0.585 | 95.451 |
| **PC20** | 0.745 | 0.532 | 95.983 |
| **PC21** | 0.668 | 0.477 | 96.460 |
| **PC22** | 0.621 | 0.444 | 96.904 |
| **PC23** | 0.567 | 0.405 | 97.309 |
| **PC24** | 0.553 | 0.395 | 97.704 |
| **PC25** | 0.470 | 0.336 | 98.040 |
| **PC26** | 0.415 | 0.297 | 98.337 |
| **PC27** | 0.332 | 0.237 | 98.574 |
| **PC28** | 0.293 | 0.209 | 98.783 |
| **PC29** | 0.250 | 0.179 | 98.962 |
| **PC30** | 0.227 | 0.162 | 99.123 |
| **PC31** | 0.194 | 0.139 | 99.262 |
| **PC32** | 0.186 | 0.133 | 99.395 |
| **PC33** | 0.172 | 0.123 | 99.518 |
| **PC34** | 0.168 | 0.120 | 99.638 |
| **PC35** | 0.142 | 0.102 | 99.740 |
| **PC36** | 0.122 | 0.087 | 99.827 |
| **PC37** | 0.085 | 0.061 | 99.888 |
| **PC38** | 0.080 | 0.057 | 99.945 |
| **PC39** | 0.077 | 0.055 | 100.000 |

**Appendix Table 1.** Full list of principal components resulting from the geometric morphometric analysis of mandibular processes. PCs 1-4 accounted for >10% of variation.

| *2x exaggerated* | Minimum Component Shape | Maximum Component Shape |
|---|---|---|
| PC1<br><br>Proportion of Variance<br>**27.23%**<br><br>Cumulative Proportion<br>**27.23%**<br><br>Eigenvalue<br>**38.13** | 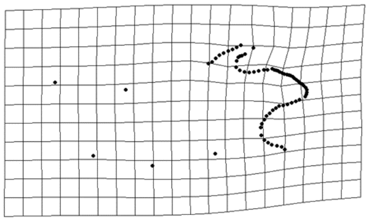 | 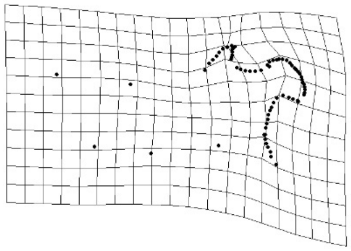 |
| PC2<br><br>Proportion of Variance<br>**17.71%**<br><br>Cumulative Proportion<br>**44.95%**<br><br>Eigenvalue<br>**24.80** | 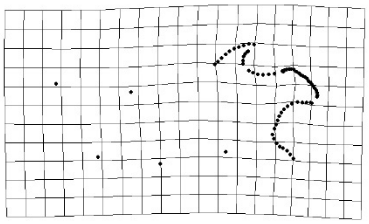 | 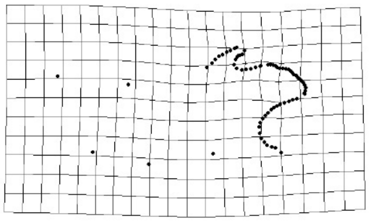 |
| PC3<br><br>Proportion of Variance<br>**13.16%**<br><br>Cumulative Proportion<br>**58.11%**<br><br>Eigenvalue<br>**18.4** | 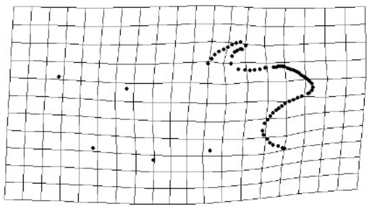 | 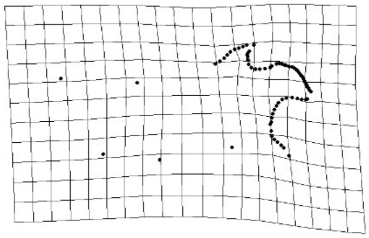 |
| PC4<br><br>Proportion of Variance<br>**10.01%**<br><br>Cumulative Proportion<br>**68.12%**<br><br>Eigenvalue<br>**14.02** | 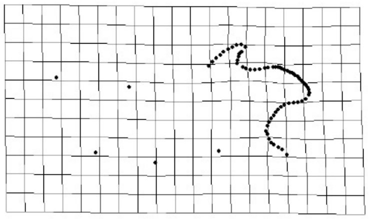 | 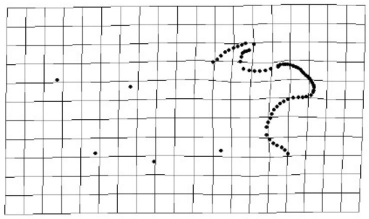 |

**Appendix Table 2**. PC1-4 were the only PCs describing >10% of variation in the geometric morphometric analysis of mandible processes. Minimum and maximum component shape are shown with 2x exaggeration for each of these PCs.

**MANCOVA Null Model**

| | df | SS | MS | Rsq | F | Z | Pr(>F) |
|---|---|---|---|---|---|---|---|
| **Centroid** | 1 | 119.20 | 119.200 | 0.032 | 1.836 | 1.120 | 0.132 |
| **Genotype** | 2 | 1147.70 | 573.840 | 0.309 | 8.839 | 5.754 | **0.0001** |
| **Residuals** | 36 | 2337.00 | 64.920 | 0.628 | | | |
| **Total** | 39 | 3719.60 | | | | | |

**Pairwise Comparison**

| | d | UCL (95%) | Z | Pr(>d) |
|---|---|---|---|---|
| **Ctrl vs GOF** | 0.029 | 0.017 | 3.658 | **0.0001** |
| **Ctrl vs LOF** | 0.017 | 0.016 | 2.021 | **0.021** |
| **GOF vs LOF** | 0.030 | 0.016 | 4.218 | **0.0001** |

**MANCOVA Interaction Model**

| | df | SS | MS | Rsq | F | Z | Pr(>F) |
|---|---|---|---|---|---|---|---|
| **Centroid** | 1 | 76.70 | 76.717 | 0.021 | 1.299 | 0.626 | 0.269 |
| **Genotype** | 2 | 329.90 | 164.931 | 0.089 | 2.792 | 2.235 | **0.011** |
| **Centroid: Genotype** | 2 | 328.50 | 164.259 | 0.088 | 2.781 | 2.226 | **0.012** |
| **Residuals** | 34 | 2008.50 | 59.074 | 0.540 | | | |
| **Total** | 39 | 3719.60 | | | | | |

**Coefficients Table**

| | d.obs | UCL (95%) | Zd | Pr(>d) |
|---|---|---|---|---|
| **Centroid** | 63.600 | 83.135 | 0.787 | 0.2244 |
| **Genotype GOF** | 1485.511 | 1042.794 | 2.960 | **0.0009** |
| **Genotype LOF** | 332.479 | 799.348 | -0.712 | 0.7583 |
| **Centroid:Genotype GOF** | 242.796 | 170.614 | 2.961 | **0.0009** |
| **Centroid:Genotype LOF** | 54.157 | 130.131 | -0.714 | 0.7582 |

**Appendix Table 3**. Statistics relating to geometric morphometric analysis of mandibular processes. MANCOVA Null Model: The effects of size and genotype are looked at separately in this model to see if they explain the overall shape variation in the first four principal components (PCs representing >10% of variation). Genotype significantly influences mandibular process morphology and more heavily influences mandibular condyle shape than centroid size. Pairwise Comparison: That genotype influences mandibular condyle shape is driven by differences between all three genotypes. MANCOVA Interaction Model: The effects of size and genotype are looked at together in this model. As in the Null Model, genotype but not centroid significantly influence mandibular process morphology. Additionally, there is a significant interaction of centroid and genotype indicating mandible size correlates with mandibular process morphology within a genotype (allometry). Coefficients Table: This allometric relationship between mandible size and mandibular process morphology occurs in the Wnt5a GOF compared to controls but not Wnt5a LOF mice compared to controls.

| | Wnt5afl/fl (Ctrl) | | | Ctsk-Cre;Wnt5afl/fl (LOF) | | | Ctsk-Cre;Rosa26-LSL-Wnt5a(GOF) | | | Unpaired t test with Welch correction, FDR (Q)=5: Wnt5afl/fl vs Ctsk-Cre;Wnt5afl/fl (LOF) | | Unpaired t test with Welch correction, FDR (Q)=5: Wnt5afl/fl vs Ctsk-Cre;Rosa26-LSL-Wnt5a (GOF) | |
|---|---|---|---|---|---|---|---|---|---|---|---|---|---|
| | **Mean** | **StDev** | **N** | **Mean** | **StDev** | **N** | **Mean** | **StDev** | **N** | **P value** | **effect size (d)** | **P value** | **effect size (d)** |
| Basisphenoid (rostral) | 0.0268 | 0.0020 | 6 | 0.0307 | 0.0019 | 6 | 0.0279 | 0.0010 | 8 | **0.0058** | **-2.018** | 0.2516 | NA |
| Basisphenoid (caudal) | 0.0481 | 0.0014 | 6 | 0.0501 | 0.0034 | 6 | 0.0493 | 0.0009 | 8 | 0.2227 | NA | 0.1055 | NA |
| Cranial base (posteriorregion) | 0.0970 | 0.0027 | 6 | 0.0996 | 0.0026 | 6 | 0.0981 | 0.0016 | 8 | 0.1165 | NA | 0.4131 | NA |
| Inter-mastoidwidth | 0.2775 | 0.0043 | 6 | 0.2828 | 0.0058 | 6 | 0.2735 | 0.0045 | 8 | 0.1027 | NA | 0.1202 | NA |
| Inter-zygomatic root width | 0.0616 | 0.0035 | 6 | 0.0623 | 0.0038 | 6 | 0.0599 | 0.0065 | 8 | 0.7702 | NA | 0.5430 | NA |
| Anterior cranial vault width | 0.1545 | 0.0069 | 6 | 0.1598 | 0.0073 | 6 | 0.1595 | 0.0040 | 8 | 0.2209 | NA | 0.1535 | NA |
| Interior frontal arch width | 0.0854 | 0.0052 | 6 | 0.0870 | 0.0054 | 6 | 0.0892 | 0.0035 | 8 | 0.5984 | NA | 0.1606 | NA |
| Inter-zygomatic arch width | 0.2013 | 0.0031 | 6 | 0.2068 | 0.0055 | 6 | 0.2061 | 0.0028 | 8 | 0.0654 | NA | 0.0130 | NA |
| Inter-orbital width | 0.1220 | 0.0052 | 6 | 0.1288 | 0.0056 | 6 | 0.1226 | 0.0037 | 8 | 0.0524 | **-1.271** | 0.8073 | NA |
| Anterior nasal width | 0.0613 | 0.0026 | 6 | 0.0639 | 0.0027 | 6 | 0.0599 | 0.0018 | 8 | 0.1288 | NA | 0.2775 | NA |
| Inter-maxillary width | 0.1267 | 0.0026 | 6 | 0.1347 | 0.0060 | 6 | 0.1275 | 0.0018 | 8 | 0.0200 | NA | 0.5151 | NA |
| Inter-molar width | 0.0916 | 0.0039 | 6 | 0.0992 | 0.0027 | 6 | 0.0896 | 0.0031 | 8 | **0.0035** | **-2.270** | 0.3345 | NA |
| Palatal width | 0.0338 | 0.0011 | 6 | 0.0356 | 0.0016 | 6 | 0.0353 | 0.0014 | 8 | 0.0425 | NA | 0.0454 | NA |
| Posterior cranial vault | 0.1327 | 0.0015 | 6 | 0.1357 | 0.0054 | 6 | 0.1376 | 0.0035 | 8 | 0.2417 | NA | **0.0053** | **-1.724** |
| Middle cranial vault | 0.1650 | 0.0032 | 6 | 0.1702 | 0.0055 | 6 | 0.1710 | 0.0040 | 8 | 0.0818 | NA | **0.0094** | **-1.613** |
| Anterior cranial vault | 0.1542 | 0.0035 | 6 | 0.1585 | 0.0051 | 6 | 0.1606 | 0.0030 | 8 | 0.1201 | NA | **0.0046** | **-2.020** |
| Frontal crest height | 0.0998 | 0.0027 | 6 | 0.1046 | 0.0034 | 6 | 0.1061 | 0.0043 | 8 | 0.0249 | NA | **0.0060** | **-1.691** |
| Anterior nasal height | 0.0486 | 0.0011 | 6 | 0.0485 | 0.0030 | 6 | 0.0483 | 0.0022 | 8 | 0.9405 | NA | 0.7596 | NA |
| Facial height | 0.0921 | 0.0029 | 6 | 0.0990 | 0.0091 | 6 | 0.0946 | 0.0044 | 8 | 0.1256 | NA | 0.2328 | NA |
| Posterior nasal height | 0.0262 | 0.0008 | 6 | 0.0276 | 0.0020 | 6 | 0.0256 | 0.0023 | 8 | 0.1743 | NA | 0.4864 | NA |
| Anterior pharyngeal height | 0.0245 | 0.0016 | 6 | 0.0236 | 0.0007 | 6 | 0.0249 | 0.0009 | 8 | 0.2325 | NA | 0.6205 | NA |
| Posterior pharyngeal height | 0.0372 | 0.0013 | 6 | 0.0362 | 0.0017 | 6 | 0.0362 | 0.0019 | 8 | 0.2757 | NA | 0.2893 | NA |
| Ear height | 0.1797 | 0.0049 | 6 | 0.1847 | 0.0061 | 6 | 0.1854 | 0.0055 | 8 | 0.1479 | NA | 0.0643 | NA |
| Palate height | 0.0176 | 0.0010 | 6 | 0.0170 | 0.0015 | 6 | 0.0155 | 0.0036 | 8 | 0.4188 | NA | 0.1478 | NA |

**Appendix Table 4.** Traditional morphometrics - means, standard deviations, and sample size**.** All measurements relative to centroid size (not raw measurements), except as noted. Statistics imported from GraphPad Prism.

| | Wnt5afl/fl (Ctrl) | | | Ctsk-Cre;Wnt5afl/fl (LOF) | | | Ctsk-Cre;Rosa26-LSL-Wnt5a(GOF) | | | Unpaired t test with Welch correction, FDR (Q)=5: Wnt5afl/fl vs Ctsk-Cre;Wnt5afl/fl (LOF) | | Unpaired t test with Welch correction, FDR (Q)=5: Wnt5afl/fl vs Ctsk-Cre;Rosa26-LSL-Wnt5a (GOF) | |
|---|---|---|---|---|---|---|---|---|---|---|---|---|---|
| | Mean | StDev | N | Mean | StDev | N | Mean | StDev | N | P value | effect size (d) | P value | effect size (d) |
| Total skull length | 0.4220 | 0.0034 | 6 | 0.4148 | 0.0051 | 6 | 0.4221 | 0.0041 | 8 | 0.0190 | **1.658** | 0.9515 | NA |
| Cranial vault length | 0.3090 | 0.0035 | 6 | 0.3155 | 0.0045 | 6 | 0.3093 | 0.0030 | 8 | 0.0203 | NA | 0.8911 | NA |
| Cranial base (rostral) | 0.1957 | 0.0016 | 6 | 0.1975 | 0.0032 | 6 | 0.1894 | 0.0023 | 8 | 0.2503 | NA | **0.0001** | **3.107** |
| Anterior cranial base | 0.0735 | 0.0028 | 6 | 0.0778 | 0.0018 | 6 | 0.0759 | 0.0025 | 8 | 0.0124 | NA | 0.1271 | NA |
| Basiocciput | 0.0794 | 0.0023 | 6 | 0.0801 | 0.0031 | 6 | 0.0760 | 0.0016 | 8 | 0.6623 | NA | 0.0119 | NA |
| Basisphenoid | 0.0566 | 0.0014 | 6 | 0.0573 | 0.0022 | 6 | 0.0568 | 0.0013 | 8 | 0.5111 | NA | 0.7868 | NA |
| Presphenoid | 0.0680 | 0.0011 | 6 | 0.0687 | 0.0010 | 6 | 0.0664 | 0.0010 | 8 | 0.2750 | NA | 0.0191 | NA |
| Facial region length | 0.2050 | 0.0046 | 6 | 0.1952 | 0.0074 | 6 | 0.2013 | 0.0038 | 8 | 0.0230 | NA | 0.1380 | NA |
| Palate length | 0.0447 | 0.0013 | 6 | 0.0443 | 0.0012 | 6 | 0.0421 | 0.0016 | 8 | 0.5267 | NA | **0.0052** | **1.786** |
| Maxilla length | 0.0923 | 0.0037 | 6 | 0.0834 | 0.0044 | 6 | 0.0878 | 0.0019 | 8 | **0.0037** | **2.185** | 0.0320 | NA |
| Premaxilla length | 0.0790 | 0.0016 | 6 | 0.0779 | 0.0027 | 6 | 0.0810 | 0.0030 | 8 | 0.4165 | NA | 0.1362 | NA |
| Nasal length | 0.1052 | 0.0050 | 6 | 0.1027 | 0.0027 | 6 | 0.1090 | 0.0024 | 8 | 0.3162 | NA | 0.1254 | NA |
| Zygomatic length | 0.0793 | 0.0019 | 6 | 0.0771 | 0.0049 | 6 | 0.0823 | 0.0028 | 8 | 0.3418 | NA | 0.0323 | NA |
| UpperJaw length | 0.2717 | 0.0023 | 6 | 0.2603 | 0.0054 | 6 | 0.2703 | 0.0056 | 8 | **0.0023** | **2.727** | 0.5354 | NA |
| Total skull length projected | 0.4210 | 0.0037 | 6 | 0.4135 | 0.0045 | 6 | 0.4214 | 0.0039 | 8 | 0.0111 | NA | 0.8574 | NA |
| Cranial vault length projected | 0.2977 | 0.0071 | 6 | 0.3023 | 0.0061 | 6 | 0.2981 | 0.0038 | 8 | 0.2500 | NA | 0.8903 | NA |
| Cranial base (rostral) projected | 0.1783 | 0.0079 | 6 | 0.1827 | 0.0033 | 6 | 0.1768 | 0.0045 | 8 | 0.2583 | NA | 0.6736 | NA |
| Anterior cranial base projected | 0.0452 | 0.0083 | 6 | 0.0446 | 0.0047 | 6 | 0.0427 | 0.0034 | 8 | 0.8913 | NA | 0.5092 | NA |
| Basiocciput projected | 0.0738 | 0.0039 | 6 | 0.0747 | 0.0020 | 6 | 0.0713 | 0.0010 | 8 | 0.6103 | NA | 0.1845 | NA |
| Basisphenoid projected | 0.0537 | 0.0027 | 6 | 0.0542 | 0.0034 | 6 | 0.0539 | 0.0025 | 8 | 0.7533 | NA | 0.8405 | NA |
| Presphenoid projected | 0.0637 | 0.0026 | 6 | 0.0645 | 0.0015 | 6 | 0.0626 | 0.0014 | 8 | 0.5425 | NA | 0.3880 | NA |
| Facial region length projected | 0.2047 | 0.0055 | 6 | 0.1950 | 0.0074 | 6 | 0.2009 | 0.0040 | 8 | 0.0300 | NA | 0.1894 | NA |
| Palate projected | 0.0399 | 0.0031 | 6 | 0.0406 | 0.0023 | 6 | 0.0382 | 0.0023 | 8 | 0.6506 | NA | 0.2912 | NA |
| Maxilla projected | 0.0901 | 0.0038 | 6 | 0.0807 | 0.0040 | 6 | 0.0851 | 0.0023 | 8 | **0.0020** | **2.397** | 0.0236 | NA |
| Premaxilla projected | 0.0750 | 0.0022 | 6 | 0.0739 | 0.0027 | 6 | 0.0777 | 0.0033 | 8 | 0.4455 | NA | 0.1000 | NA |

**Appendix Table 4 (continued).** Traditional morphometrics - means, standard deviations, and sample size**.** All measurements relative to centroid size (not raw measurements), except as noted. Statistics imported from GraphPad Prism.

| | Wnt5afl/fl (Ctrl) | | | Ctsk-Cre;Wnt5afl/fl (LOF) | | | Ctsk-Cre;Rosa26-LSL-Wnt5a(GOF) | | | Unpaired t test with Welch correction, FDR (Q)=5 | | | |
|---|---|---|---|---|---|---|---|---|---|---|---|---|---|
| | | | | | | | | | | Wnt5afl/fl vs Ctsk-Cre;Wnt5afl/fl (LOF) | | Wnt5afl/fl vs Ctsk-Cre;Rosa26-LSL-Wnt5a (GOF) | |
| | Mean | StDev | N | Mean | StDev | N | Mean | StDev | N | P value | effect size (d) | P value | effect size (d) |
| Nasal projected | 0.0967 | 0.0047 | 6 | 0.0908 | 0.0061 | 6 | 0.0996 | 0.0034 | 8 | 0.0907 | NA | 0.2268 | NA |
| Zygomatic projected | 0.0755 | 0.0015 | 6 | 0.0742 | 0.0045 | 6 | 0.0796 | 0.0029 | 8 | 0.5295 | NA | **0.0060** | **-1.677** |
| UpperJaw projected | 0.2432 | 0.0035 | 6 | 0.2310 | 0.0060 | 6 | 0.2400 | 0.0062 | 8 | **0.0027** | **2.459** | 0.2541 | NA |
| Mandibular posterior height | 0.0551 | 0.0014 | 6 | 0.0552 | 0.0020 | 6 | 0.0553 | 0.0023 | 8 | 0.9354 | NA | 0.8149 | NA |
| Mandibularlength (superior) | 0.2317 | 0.0023 | 6 | 0.2305 | 0.0021 | 6 | 0.2235 | 0.0015 | 8 | 0.3823 | NA | **0.0001** | **4.298** |
| Mandibularlength (inferior) | 0.1745 | 0.0010 | 6 | 0.1733 | 0.0031 | 6 | 0.1669 | 0.0014 | 8 | 0.4124 | NA | **<0.000001** | **6.162** |
| Inter-molar width (mandible) | 0.0835 | 0.0033 | 6 | 0.0841 | 0.0033 | 6 | 0.0880 | 0.0026 | 8 | 0.7476 | NA | 0.0231 | NA |
| Bi-condylar width | 0.2247 | 0.0039 | 6 | 0.2317 | 0.0039 | 6 | 0.2244 | 0.0030 | 8 | 0.0116 | NA | 0.8831 | NA |
| Bi-gonial width | 0.1868 | 0.0044 | 6 | 0.1930 | 0.0044 | 6 | 0.1950 | 0.0027 | 8 | 0.0368 | NA | **0.0043** | **-2.303** |
| Mandibular anterior height | 0.0413 | 0.0085 | 6 | 0.0346 | 0.0028 | 6 | 0.0393 | 0.0013 | 8 | 0.1154 | NA | 0.5908 | NA |
| Condylar width | 0.0393 | 0.0027 | 6 | 0.0405 | 0.0015 | 6 | 0.0410 | 0.0011 | 8 | 0.3512 | NA | 0.1814 | NA |
| Angular Process superior length | 0.0292 | 0.0013 | 6 | 0.0308 | 0.0017 | 6 | 0.0296 | 0.0007 | 8 | 0.0971 | NA | 0.5456 | NA |
| Angular Process inferior length | 0.0578 | 0.0033 | 6 | 0.0537 | 0.0036 | 6 | 0.0509 | 0.0035 | 8 | 0.0643 | NA | **0.0028** | **2.049** |
| coronoid-condylar inlet | 0.0003 | 0.0000 | 6 | 0.0003 | 0.0001 | 6 | 0.0002 | 0.0000 | 8 | 0.0817 | NA | **0.0007** | **2.661** |
| posterior mandibular inlet | 0.0007 | 0.0000 | 6 | 0.0008 | 0.0000 | 6 | 0.0007 | 0.0000 | 8 | 0.4880 | NA | 0.0176 | NA |
| ventral mandibular inlet | 0.0006 | 0.0000 | 6 | 0.0007 | 0.0000 | 6 | 0.0007 | 0.0000 | 8 | 0.0816 | NA | **0.0004** | **-2.873** |
| inter-ear width upper | 0.2244 | 0.0031 | 6 | 0.2308 | 0.0024 | 6 | 0.2288 | 0.0037 | 8 | **0.0027** | **-2.322** | 0.0328 | NA |
| inter-ear width lower | 0.1703 | 0.0117 | 6 | 0.1824 | 0.0049 | 6 | 0.1684 | 0.0116 | 8 | 0.0546 | NA | 0.7691 | NA |
| inter-ear width average | 0.1973 | 0.0063 | 6 | 0.2066 | 0.0025 | 6 | 0.1986 | 0.0060 | 8 | 0.0138 | NA | 0.7199 | NA |
| RAW Maxillary molar field length | 2.7833 | 0.0745 | 6 | 2.5700 | 0.1621 | 6 | 2.8275 | 0.1540 | 8 | 0.0220 | **1.691** | 0.4940 | NA |
| RAW Mandibular molar field length | 3.2817 | 0.0711 | 6 | 3.0683 | 0.1446 | 6 | 3.2013 | 0.0710 | 8 | 0.0134 | **1.873** | 0.0603 | NA |

**Appendix Table 4 (continued).** Traditional morphometrics - means, standard deviations, and sample size**.** All measurements relative to centroid size (not raw measurements), except as noted. Statistics imported from GraphPad Prism.

| Measurement | %CV | Type |
|---|---|---|
| Basisphenoid (rostral) | 1.64 | linear |
| Basisphenoid (caudal) | 2.21 | linear |
| Cranial base (posteriorregion) | 0.32 | linear |
| Inter-mastoidwidth | 0.30 | linear |
| Inter-zygomatic root width R | 0.59 | linear |
| Inter-zygomatic root width L | 0.34 | linear |
| Inter-zygomatic root width Avg | 0.47 | linear |
| Anterior cranial vault width | 0.37 | linear |
| Inter-ear width R | 0.42 | linear |
| Inter-ear width L | 4.14 | linear |
| Inter-ear width Avg | 2.01 | linear |
| Interior frontal arch width | 3.36 | linear |
| Inter-zygomatic arch width | 0.55 | linear |
| Inter-orbital width | 0.42 | linear |
| Anterior nasal width | 0.38 | linear |
| Inter-maxillary width | 1.33 | linear |
| Inter-molar width | 0.02 | linear |
| Palatal width | 0.07 | linear |
| Posterior cranial vault | 0.01 | linear |
| Middle cranial vault | 0.02 | linear |
| Anterior cranial vault | 0.30 | linear |
| Frontal crest height | 0.14 | linear |
| Anterior nasal height | 0.80 | linear |
| Facial height | 0.13 | linear |
| Posterior nasal height | 0.15 | linear |
| Anterior pharyngeal height | 1.88 | linear |
| Posterior pharyngeal height R | 0.01 | linear |
| Posterior pharyngeal height L | 0.26 | linear |
| Posterior pharyngeal height Avg | 0.13 | linear |
| Ear height R up | 0.74 | linear |
| Ear height R down | 0.90 | linear |
| Ear height R Avg | 0.83 | linear |
| Ear height L up | 0.54 | linear |
| Ear height L down | 0.56 | linear |
| Ear height L Avg | 0.09 | linear |
| Ear height Avg | 0.46 | linear |
| Total skull length | 0.19 | linear |
| Cranial vault length | 0.24 | linear |
| Cranial base (rostral) | 0.02 | linear |
| Anterior cranial base | 0.21 | linear |
| Basiocciput | 0.09 | linear |
| Basisphenoid | 0.52 | linear |
| Presphenoid | 0.16 | linear |
| Facial region length | 0.12 | linear |
| Palate | 0.04 | linear |
| Maxilla R | 0.23 | linear |
| Maxilla L | 0.13 | linear |
| Maxilla Avg | 0.05 | linear |
| Premaxilla R | 0.56 | linear |
| Premaxilla L | 0.13 | linear |
| Premaxilla Avg | 0.35 | linear |
| Nasal | 1.51 | linear |
| Zygomatic R | 0.31 | linear |
| Zygomatic L | 0.38 | linear |
| Zygomatic Avg | 0.35 | linear |
| UpperJaw R front | 0.66 | linear |

| Measurement | %CV | Type |
|---|---|---|
| UpperJaw R back | 0.78 | linear |
| UpperJaw R Avg | 0.73 | linear |
| UpperJaw L front | 0.84 | linear |
| UpperJaw L back | 0.84 | linear |
| UpperJaw L Avg | 0.84 | linear |
| UpperJaw Avg | 0.78 | linear |
| Mandibular posterior height R | 0.62 | linear |
| Mandibular posterior height L | 0.36 | linear |
| Mandibular posterior height Avg | 0.49 | linear |
| Mandibularlength (superior) R | 0.12 | linear |
| Mandibularlength (superior) L | 0.00 | linear |
| Mandibularlength (superior) Avg | 0.06 | linear |
| Mandibularlength (inferior) R | 0.05 | linear |
| Mandibularlength (inferior) L | 0.01 | linear |
| Mandibularlength (inferior) Avg | 0.02 | linear |
| Mandibular inter-molar width | 2.08 | linear |
| Bi-condylar width | 0.29 | linear |
| Bi-gonial width | 0.03 | linear |
| Mandibular anterior height R | 0.07 | linear |
| Mandibular anterior height L | 1.86 | linear |
| Mandibular anterior height Avg | 0.95 | linear |
| Condylar width R | 2.62 | linear |
| Condylar width L | 2.94 | linear |
| Condylar width Avg | 2.78 | linear |
| Angular Process superior length R | 0.92 | linear |
| Angular Process superior length L | 1.54 | linear |
| Angular Process superior length Avg | 0.31 | linear |
| Angular Process inferior length R | 2.43 | linear |
| Angular Process inferior length L | 3.05 | linear |
| Angular Process inferior length Avg | 2.74 | linear |
| Coronoid-Condylar inlet R | 3.50 | angular |
| Coronoid-Condylar inlet L | 6.09 | angular |
| Coronoid-Condylar inlet Avg | 4.77 | angular |
| Posterior mandibular inlet R | 0.36 | angular |
| Posterior mandibular inlet L | 1.94 | angular |
| Posterior mandibular inlet Avg | 1.16 | angular |
| Ventral mandibular inlet R | 1.69 | angular |
| Ventral mandibular inlet L | 0.19 | angular |
| Ventral mandibular inlet Avg | 0.73 | angular |
| Cranial base angle | 0.09 | angular |
| Anterior cranial vault angle | 0.15 | angular |
| Mid-anterior cranial vault angle | 0.43 | angular |
| Mid-posterior cranial vault angle | 0.15 | angular |
| Posterior cranial vault angle | 0.14 | angular |
| Snout angle | 1.69 | angular |
| Facial angle | 0.29 | angular |
| Palatte arch angle | 0.40 | angular |
| Nasale-Lambda midline nasal deviation | 0.87 | angular |
| Bregma-Lambda midline nasal deviation | 1.00 | angular |
| Face tipping angle | 0.13 | angular |
| Mid-saggital plane midline nasal deviation | 9.32 | angular |
| Cranial Centroid Size (mm) | 0.09 | neither |
| Mandibular Centroid Size (mm) | 0.16 | neither |
| L-Hemimandible Centroid Size (mm) | 0.18 | neither |
| R-Hemimandible Centroid Size (mm) | 0.14 | neither |

**Appendix Table 5**. Intraobserver percent coefficient of variation (CV) values. As is standard, a percent CV value of ≤10% was deemed acceptable replicability. R=right; L=left, Avg=average

| Summary | | *Wnt5a* LOF *Wnt5a-flox; Ctsk-cre* | *Wnt5a* GOF *Rosa26-LSL-Wnt5a;Ctsk-cre* | Correlating reported clinical phenotype |
|---|---|---|---|---|
| | body weight | ↓P70 ♂ | --- | |
| **craniofacial bone** (this paper) | maxilla length | ↓ | --- | midface hypoplasia |
| | upper jaw length | ↓ | --- | midface hypoplasia |
| | maxillary intermolar width | ↑ | --- | |
| | rostral basisphenoid width | ↑ | --- | |
| | tooth eruption | delayed | --- | *** |
| | snout deviation | right sided | right sided | *** |
| | mandible length | --- | ↓ | micrognathia |
| | mandible bone surface | --- | ↓ | micrognathia |
| | mandibular angular process length | --- | ↓ | |
| | coronoid and condylar processes morphology | --- | morphology Δ | *** |
| | cranial vault height | --- | ↑ | macrocephaly |
| | palate length | --- | ↓ | short hard palate |
| | zygomatic length | --- | ↑ | |
| | mandible trabecular connectivity | --- | ↓ | |
| | mandible bone surface | --- | ↓ | micrognathia |
| **long bone** (Roberts et al. 2020) | femur BMD | ↓ | not assessed | non-cranial osteopenia |
| | femur trabecular thickness | ↓ | not assessed | |
| | femur trabecular separation | ↑ | not assessed | |
| | femur bone surface | ↓ | not assessed | |

**Appendix Table 6.** Summary of key findings. Up or down arrow indicates statistically significant increase or decreased in the measure in the indicated group compared to controls, dash indicates no difference. *** should be evaluated in patients with Robinow Syndrome.
